# Cellular and Molecular Architecture of Renin-Angiotensin System Signaling in the PVN Under Cardiometabolic Stress

**DOI:** 10.1101/2025.08.14.669892

**Authors:** Haifeng Zheng, Ha Nguyen, Khoi Nguyen, Shiyue Pan, Tong Zhou, Tin Nguyen, Yumei Feng Earley

**Affiliations:** Department of Medicine, University of Rochester Medical Center, Rochester, NY, 14642, USA; Department of Pharmacology & Physiology, University of Rochester Medical Center, Rochester, NY, 14642, USA; Department of Computer Science and Software Engineering, Auburn University, Auburn, AL, 36849, USA; Department of Physiology and Cell Biology, University of Nevada, Reno, NV 89557 USA

**Keywords:** renin-angiotensin system, hypothalamic paraventricular nucleus, single- nucleus RNA sequencing, high-fat diet–induced obesity, DOCA-salt–induced hypertension

## Abstract

The hypothalamic paraventricular nucleus (PVN) integrates neuroendocrine and autonomic signals that regulate blood pressure and metabolism. Although the renin-angiotensin system (RAS) is implicated in neurogenic hypertension and obesity, cell-type-specific expression and regulation of its components within the PVN remain poorly understood. Here, we employed single-nucleus RNA sequencing (snRNA-seq) to profile the transcriptomic landscape of the PVN in male mice under baseline conditions and in models of DOCA-salt–induced hypertension and high-fat diet (HFD)-induced obesity.

We identified major PVN cell types, including neurons, astrocytes, precursor oligodendrocytes, oligodendrocytes, microglia and endothelial cells, and further resolved eight transcriptionally distinct neuronal subtypes. Expression of RAS-related genes was highly cell-type specific: *Agt* (angiotensinogen) was enriched in astrocytes, whereas *Ace* (angiotensin-converting enzyme), *Atp6ap2* (also known as the (pro)renin receptor [PRR]), *Agtr1a* (angiotensin II type 1a receptor, aka AT^1a^R), *Lnpep* (leucyl/cystinyl aminopeptidase, aka angiotensin 4 receptor [AT^4^R]), and the *Mas1* proto-oncogene were predominantly expressed in neurons. DOCA-salt treatment increased the proportion of GABAergic and vasopressin neurons and enhanced neuronal *Agt* and *Atp6ap2* expression, while reducing astrocytic *Agt*, suggesting activation of a vasoconstrictive RAS axis. HFD exposure increased excitatory and stress-responsive neuronal subtypes (glutamatergic, vasopressin, corticotropin-releasing hormone) and upregulated *Atp6ap2*, *Agtr1b*, *Lnpep*, and *Mas1* in vasopressin neurons, while downregulating multiple RAS genes in GABAergic neurons.

These findings reveal dynamic, cell-type–specific remodeling of RAS signaling in the PVN in response to hypertensive and metabolic stress, providing a transcriptomic atlas of RAS expression in the PVN and identifying potential cellular targets for therapeutic strategies addressing cardiometabolic disorders.

## Introduction

The hypothalamic paraventricular nucleus (PVN) serves as a central integrator of neuroendocrine, autonomic, and behavioral responses critical for maintaining cardiovascular and metabolic homeostasis^1–4^. Within this region, neuronal and glial cell types tightly regulate the output of neuropeptides and autonomic signals that influence blood pressure, fluid balance, energy metabolism, and stress responses^5–7^. Disruptions in PVN function are increasingly implicated in cardiometabolic disorders, including hypertension and obesity, both of which are major global health concerns^8, 9^. One key signaling cascade that contributes to PVN-mediated physiological regulation is the brain renin-angiotensin system (RAS)^10–13^. Traditionally known for their roles in the periphery (e.g., cardiovascular and fluid homeostasis), RAS components are also expressed in distinct brain regions, including the hypothalamus^14, 15^. Central RAS activity modulates sympathetic outflow, vasopressin release, and feeding behavior^2, 12^. Notably, alterations in brain RAS signaling have been linked to the development and progression of neurogenic hypertension and metabolic dysfunction^16–20^. Despite this, the precise cellular sources and subtype-specific expression patterns of RAS components within the PVN remain incompletely understood.

The recent advent of single-cell transcriptomic technologies has revolutionized our ability to interrogate cellular heterogeneity and gene expression dynamics within complex brain structures. Among these techniques, single-nucleus RNA sequencing (snRNA-seq) offers a unique advantage, enabling the use of frozen tissues and minimizing dissociation-induced transcriptional artifacts^21–23^. While snRNA-seq captures only nuclear transcripts and may underrepresent cytoplasmic mRNAs, it provides high-resolution insight into the transcriptional landscape of cells in their native tissue architecture^21, 22^. This approach is particularly well-suited for brain tissue, which is often challenging to dissociate into viable single cells without altering gene expression profiles.

In this study, we utilized snRNA-seq to generate a comprehensive cellular and transcriptomic atlas of the PVN, with a specific focus on the expression of RAS genes across different cell types and neuronal subpopulations. We further examined how two pathophysiological models—deoxycorticosterone acetate (DOCA)-salt–induced hypertension and high-fat diet (HFD)–induced hypertension and obesity—alter PVN cellular composition and RAS gene expression. Our findings reveal distinct and cell-type– specific remodeling of the RAS axis under hypertensive and obesogenic conditions, highlighting novel mechanisms through which neurogenic RAS signaling may contribute to cardiometabolic disease. These insights lay the groundwork for future functional studies aimed at dissecting the roles of individual PVN cell types in RAS-mediated physiological regulation and may inform new therapeutic targets for hypertension and obesity.

## Materials and Methods

### Animals

C57BL/6J mice were purchased from The Jackson Laboratory (Stock #000664; Bar Harbor, ME). All animal care procedures and experimental protocols involving animals complied with the NIH Guide for the Care and Use of Laboratory Animals and were approved by the University Committee on Animal Resources (UCAR) at the University of Rochester, Rochester, NY. Mice were maintained in individually ventilated cages (<5 mice/cage) with ad libitum access to food and water in a room with controlled 12-hour light and dark cycles.

### High-Fat Diets and DOCA-Salt Treatment Protocols

Male C57BL/6J mice (12 weeks old, n = 10 mice/group) were fed ad libitum a high-fat diet (HFD, #D12492; Open Source Diets, Research Diets Inc., New Brunswick, NJ), providing 60% of kCal from fat, or a control chow diet (CD, #2019; ENVIGO, Indianapolis, IN), providing 22% of kCal from fat, for 6 weeks ^24, 25^. The DOCA-salt hypertension model was established following previously described protocols^13, 26, 27^. Briefly, C57BL/6J mice (12 weeks old, n = 10 mice/group) were randomized into two groups—a SHAM group and a DOCA-salt group—and subcutaneously implanted with a sham pellet (C-111) or 50 mg DOCA pellet (M-121; Sigma-Aldrich, St. Louis, MO, USA), respectively, below the scapula under isoflurane anesthesia. Mice were matched for body mass and composition prior to surgery. Mice in the SHAM group were given free access to regular tap water, while mice in the DOCA-salt treatment group received 0.9% saline as drinking water. Regular chow was provided ad libitum to both groups for 14 days.

### Brain Tissue Isolation, Single-Nucleus RNA

Mice were euthanized and their whole brains were dissected, after which the PVN was isolated using mouse stainless steel brain matrices (1 mm, #15003; Ted Pella, Inc.) and razors. Tissues were cut approximately from bregma -0.46 mm to -1.46 mm; hypothalamic tissue containing the PVN was dissected and flash frozen in liquid nitrogen and stored briefly (days) in a -80°C freezer during preparation for shipping. Isolation of nuclei, single-nucleus RNA sequencing, and ATAC sequencing were conducted by Singulomics Corporation (Bronx, NY). Nuclei were isolated and single-nuclei gene expression and ATAC libraries were prepared using the 10x Genomics Chromium System with the Chromium Multiome Kit (10x Genomics, Pleasanton, CA). Each library was sequenced to a depth of approximately 200 million paired-end reads (PE150) on an Illumina NovaSeq 6000 (Illumina, San Diego, CA). A summary of the protocol is presented in **Figure 1**.

**Figure 1.**
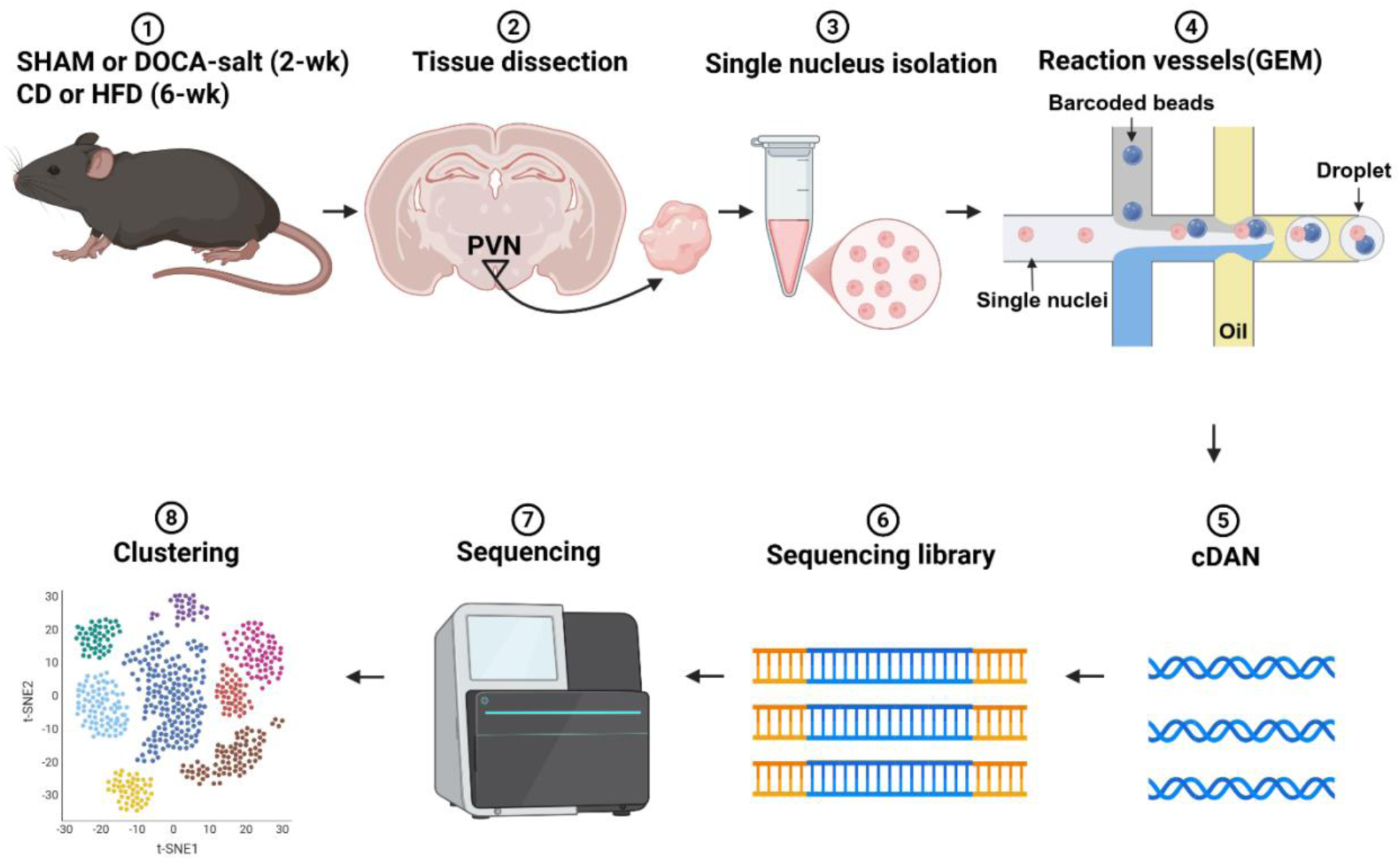
Overview of experimental workflow for snRNA-seq analysis of the PVN. Illustration of the step-by-step procedure: ① Mice underwent either SHAM or DOCA-salt treatment for 2 weeks or were fed a CD or HFD for 6 weeks. ② Brain tissue containing the PVN was carefully dissected. ③ Single-nucleus isolation was performed to obtain individualized cells from dissected PVN tissues. ④ Single nuclei were encapsulated with barcoded beads in droplets to generate Gel Beads-in-Emulsion (GEMs). ⑤ Reverse transcription within GEMs produced cDNA. ⑥ cDNA libraries were prepared and indexed for sequencing. ⑦ Sequencing was performed using a next-generation sequencing platform. ⑧ Data analysis was carried out, including clustering of cell populations using dimensionality reduction techniques (e.g., t-SNE), enabling the identification of distinct cellular subpopulations within the PVN. Figure created using BioRender.

### Single-nucleus Data Analysis Using Cell Ranger ARC

For each experiment, single-nucleus multiome data were processed using Cell Ranger ARC version 2.0.2 (10x Genomics). Raw sequencing reads in FASTQ format were aligned to the *mm10* mouse reference genome (*mm10-2020-A-2.0.0* version), and filtered feature-barcode matrices were generated for both RNA and ATAC modalities.

Low-quality barcodes and empty droplets were excluded by performing barcode selection and initial quality control with Cell Ranger ARC using default internal heuristics. The Cell Ranger ARC pipeline produced several key output files, including (1) a *web_summary.html* report, summarizing sequencing and quality control metrics for both modalities; (2) filtered and raw feature-barcode matrices in Matrix Market Exchange (MEX) format; (3) aligned BAM files and corresponding index files for both RNA and ATAC reads; and (4) ATAC peak annotation, with detailed molecule-level and other information.

### Cell-Type Annotation Using scRNA-seq Data and Seurat

Seurat objects were constructed from the filtered RNA-Seq count matrix output with Cell Ranger ARC and analyzed using Seurat v5.3.0^28–31^. Normalization and variance stabilization were performed using *SCTransform*^32^ with default parameters.

Dimensionality reduction was conducted via principal component analysis33 using the *RunPCA* function, and the top 30 principal components (PCs) were used to compute two-dimensional embeddings via Uniform Manifold Approximation and Projection (UMAP)34 and t-distributed Stochastic Neighbor Embedding (t-SNE)35 using the *RunUMAP* and *RunTSNE* functions, respectively.

Cell-type annotation was carried out in two phases. In the first phase, reference-based label transfer was performed using the Azimuth Mouse Motor Cortex reference dataset^36^. Transfer anchors (corresponding cell pairs) were identified between our dataset and the reference dataset using the *FindTransferAnchors* function on the top 10 PCs, and cell-type labels were assigned using *MapQuery*. This yielded six major cell type annotations: oligodendrocytes, precursor oligodendrocytes, astrocytes, endothelial cells, microglia, and neurons (**Table 1**).

**Table 1.**
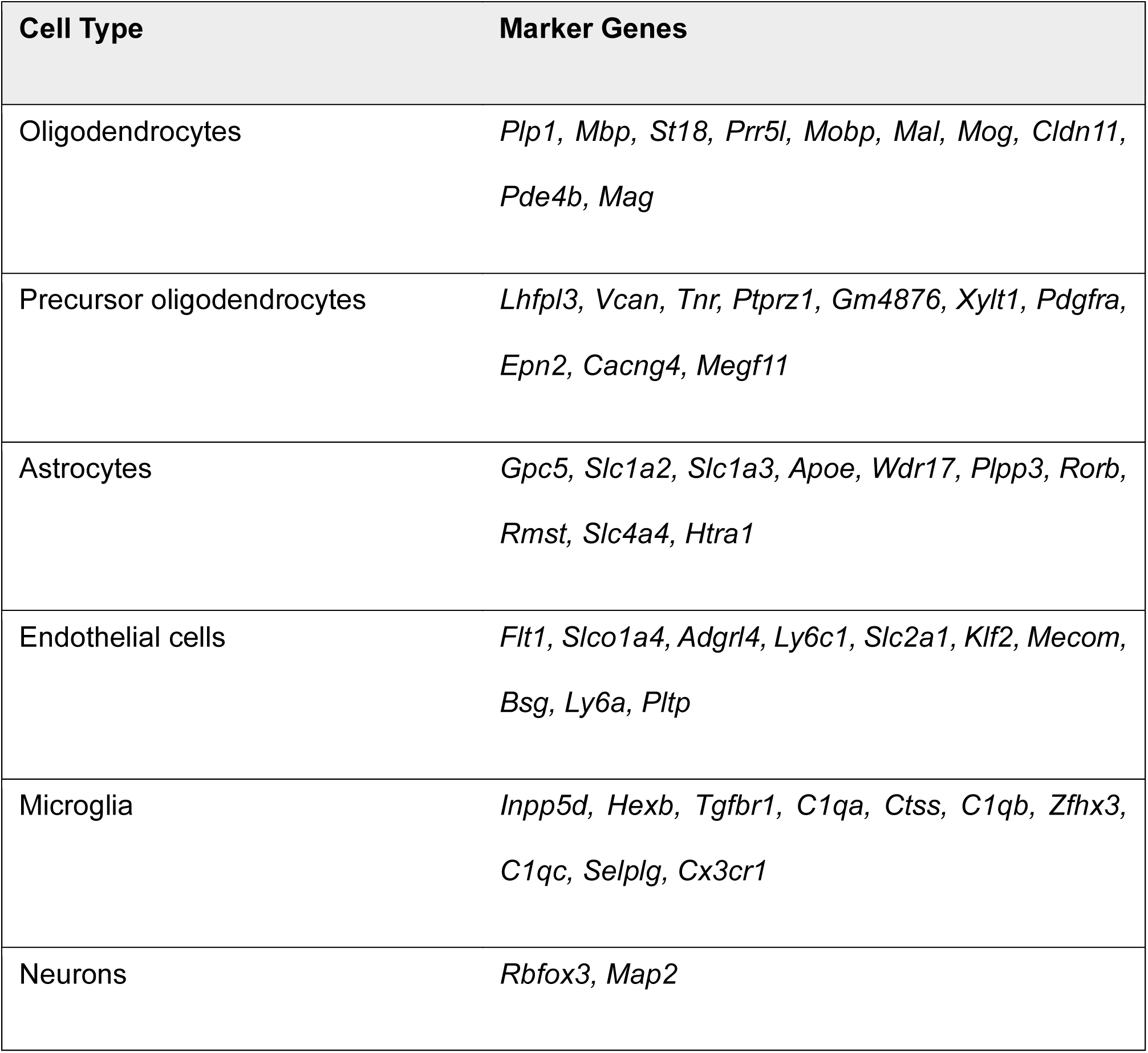
Annotated Marker Genes for Cell Type Identification.

In the second phase, cells initially annotated as neurons were further classified into subtypes based on the non-zero expression of canonical marker genes (listed in Table 2). These subtypes included glutamatergic neurons, GABAergic neurons, glutamatergic + GABAergic neurons, cholinergic neurons, TH (tyrosine hydroxylase) neurons, vasopressin neurons, CRH (corticotropin-releasing hormone) neurons, and TRH (thyrotropin-releasing hormone) neurons.

**Table 2.** Annotated Marker Genes for Neuronal Subpopulation Identification.

| **Cell Type** | **Marker Genes** |
| --- | --- |
| Glutamatergic neurons | *Rbfox3, Map2, Slc17a6, Slc17a7, Grin1* |
| GABAergic neurons | *Rbfox3, Map2, Slc32a1, Gad2* |
| Glutamatergic +<br>GABAergic neurons | *Rbfox3, Map2, Slc32a1, Gad2, Slc17a6, Slc17a7, Grin1* |
| Cholinergic neurons | *Rbfox3, Map2, Ache, Chat* |
| TH neurons | *Rbfox3, Map2, Th* |
| Vasopressin neurons | *Rbfox3, Map2, Avp* |
| CRH neurons | *Rbfox3, Map2, Crh* |
| TRH neurons | *Rbfox3, Map2, Trh* |

### Statistical Analysis

All statistical analyses were performed using Fisher’s exact test in GraphPad Prism 10 (GraphPad Software, La Jolla, CA, USA). Differences were considered statistically significant at *P*-values < 0.05.

## Results

### snRNA-seq Reveals the Cellular Composition and Transcriptomic Landscape of the RAS in the PVN

To characterize the cellular composition of the hypothalamus, we performed snRNA-seq and clustered cells based on their transcriptomic profiles. A total of 3670 cells were sequenced from mice fed a regular chow control diet (CD). A t-distributed Stochastic Neighbor Embedding (t-SNE) plot (**Figure 2A**) demonstrated distinct clustering of neuronal and non-neuronal cell populations into major cell types, including neurons, astrocytes, oligodendrocytes, microglia, precursor oligodendrocytes and endothelial cells, as well as other minor populations. Each cluster was assigned to a specific cell type following the first phase of the cell type annotation pipeline described in the Data Analysis section. A pie chart illustration of the proportional distribution of cell types (**Figure 2B**) revealed that neurons represent the largest population (45.1%), followed by oligodendrocytes (34.4%) and astrocytes (11.7%). Other glial and vascular cell types, including microglia, precursor oligodendrocytes and endothelial cells, contributed to the remaining cell fractions.

**Figure 2.**
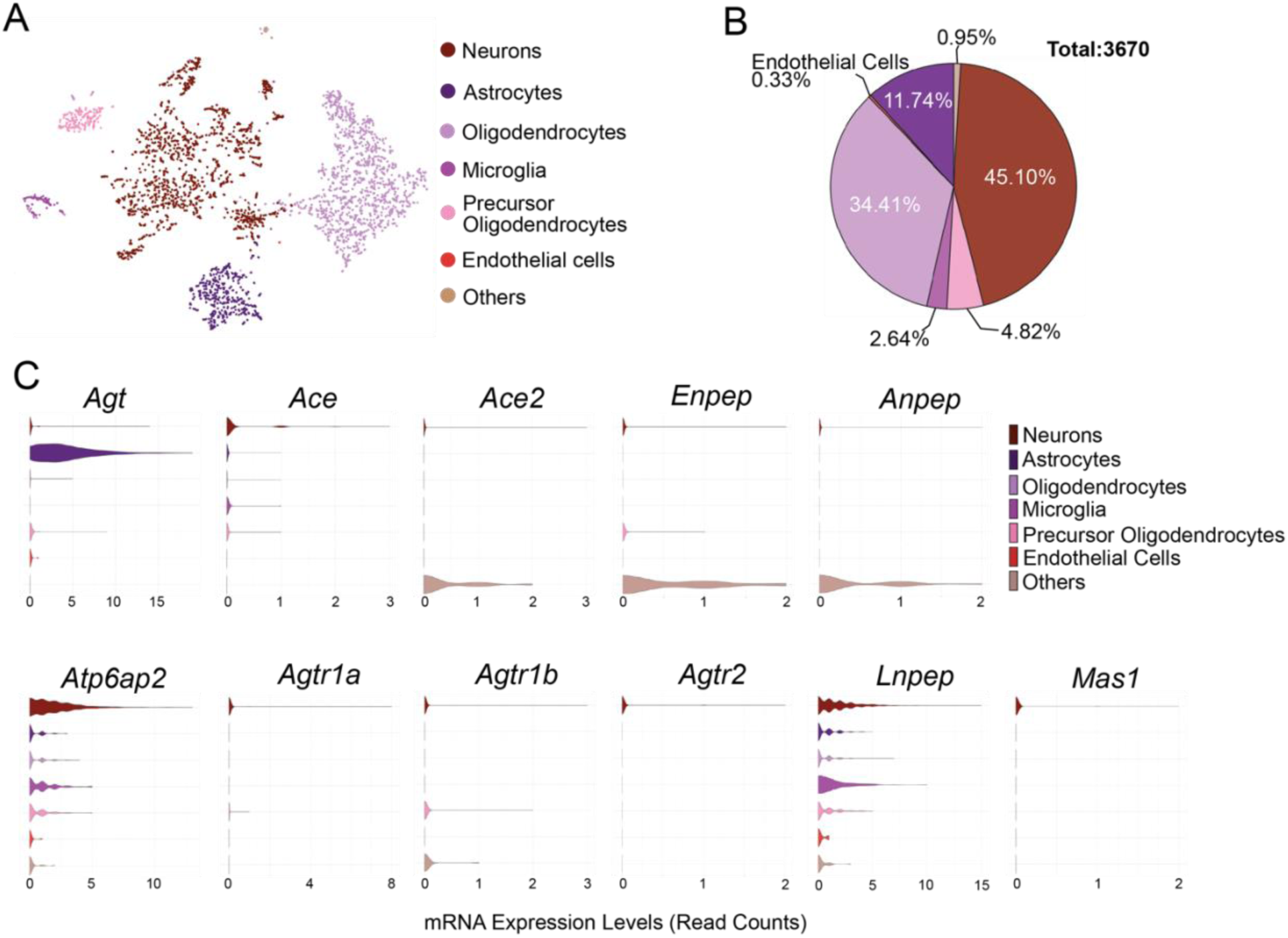
Cellular Composition and Transcriptomic Profiles of the RAS in the PVN Revealed by snRNA-seq. (A) t-SNE visualization demonstrating distinct clustering of major cell types identified in the PVN from mice fed a CD. (B) Pie chart showing the proportional distribution of major cell populations within the PVN. Numbers represent the total cell count for each population. (C) Violin plots illustrating the expression distribution of key RAS genes across different PVN cell types. Cell types are color-coded according to the legend provided.

To further define the expression pattern of RAS genes in each cell type, we analyzed their expression patterns across all identified clusters. Violin plots (**Figure 2C**) illustrate the distribution of RAS components’ expression in neurons, astrocytes, oligodendrocytes, microglia, precursor oligodendrocytes, and endothelial cells. Notably, *Agt* (angiotensinogen) was expressed at the highest levels in astrocytes, but was also expressed at lower levels in neurons and precursor oligodendrocytes. Other key RAS-related genes, including *Ace* (angiotensin-converting enzyme), *Enpep* (glutamyl aminopeptidase, also called aminopeptidase A (APA]), *Atp6ap2* (ATPase H^+^ transporting lysosomal accessory protein 2, also known as the (pro)renin receptor [PRR]), angiotensin II receptor type 1a (*Agtr1a*, also called AT^1a^R) and type 1b (*Agtr1b*, also called AT^1b^R), angiotensin II receptor type 2 (*Agtr2*, also called AT^2^R), *Lnpep* (leucyl and cystinyl aminopeptidase, also known as angiotensin IV receptor [AT^4^R]), and *Mas1* (encoding a G protein-coupled receptor oncogene) were expressed in neurons. Interestingly, *Atp6ap2* and *Lnpep* exhibited broader expression patterns in cell types beyond neurons compared with other RAS genes. Together, these findings establish a comprehensive atlas of RAS-related genes in hypothalamic PVN cells, revealing the heterogeneity of neuron and glial populations and providing insights into their unique RAS transcriptional landscapes.

### Neuronal Subtype-Specific Expression of RAS Genes in the PVN

To investigate the expression patterns of RAS-related genes in neuronal subtypes of the PVN, we performed clustering of neuronal populations based on their transcriptional profiles. A t-SNE plot (**Figure 3A**) illustrates the separation of cells into major neuronal subtypes, including TH neurons, cholinergic neurons, CRH neurons, glutamatergic and GABAergic hybrid neurons, TRH neurons, vasopressin neurons, and distinct populations of glutamatergic and GABAergic neurons. Each cluster was identified based on the expression of well-established marker genes (**Table 2**), as described in the second phase of the cell-type annotation pipeline in the Data Analysis section. The proportional distribution of neuronal subtypes is represented in a pie chart (**Figure 3B**), which demonstrates that GABAergic neurons form the predominant population (54.4%) within the PVN area. Glutamatergic neurons represent the second-most abundant group (21.8%), followed by vasopressin neurons (13.5%). Other neuronal subtypes, including cholinergic, CRH, TRH and TH-expressing neurons, accounted for smaller fractions of the total neuronal population. Violin plots (**Figure 3C**) illustrate the differential expression of key RAS genes across neuronal subtypes. *Ace*, *Atp6ap2*, *Agtr1a*, and *Lnpep* were highly expressed across all neuron types. Ace2, Enpep, Anpep, *Agtr1b,and Agtr2 expression levels were low across different neuronal types,* whereas *Mas1* was absent in TH and CRH neurons. These findings highlight the diversity of neuronal subtypes within the PVN and their distinct RAS gene expression profiles and provide a genetic basis for establishing the potential contribution of these subtypes to RAS-mediated physiological processes.

**Figure 3.**
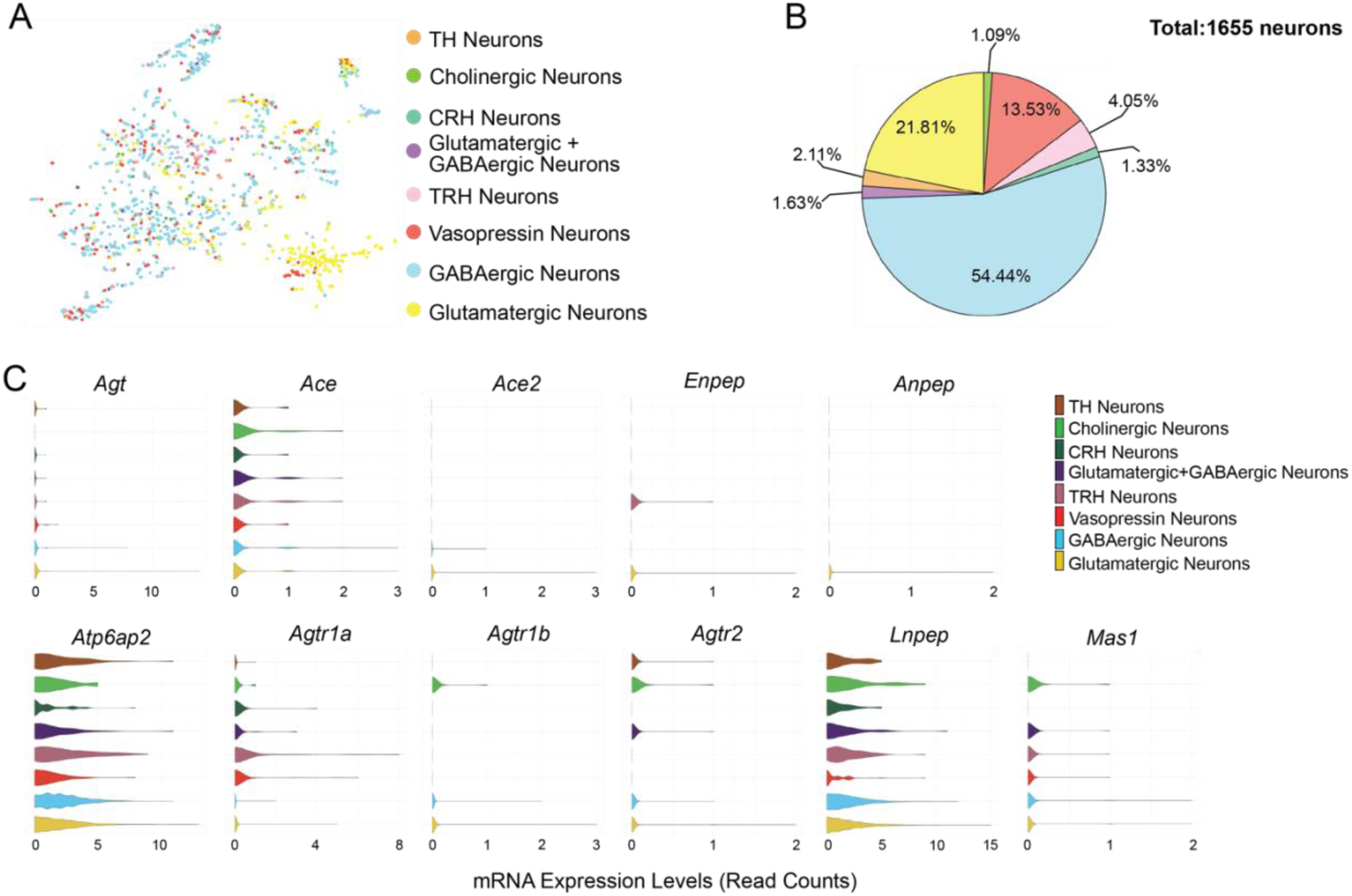
Neuronal subtype-specific expression patterns of RAS-related genes in the PVN. (A) t-SNE visualization illustrating the clustering of distinct neuronal subtypes based on their transcriptomic profiles. (B) Pie chart displaying the proportional distribution of neuronal subtypes within the PVN. (C) Violin plots demonstrating the differential expression of RAS genes across identified neuronal subtypes. Colors correspond to neuronal subtypes, as indicated in the legend.

### DOCA-salt Treatment Alters Hydration, Activity, and Sleep Behavior Without Affecting Body Weight or Energy Expenditure

Hypertension development induced by DOCA-salt has been well characterized previously ^13, 26, 27, 37^, but the impact of DOCA-salt on metabolic phenotypes has received less research attention. To evaluate the metabolic and behavioral impact of DOCA-salt– induced hypertension, we monitored mice in metabolic cages over a 60-hour period. Although DOCA-salt–treated mice exhibited no difference in food intake compared with SHAM controls (**Figure 4A**), they showed significantly increased cumulative fluid intake (**Figure 4B**) and increased water vapor (VH₂O) loss, both in circadian profiles and as averaged values across dark and light phases (**Figure 4C, D**), suggesting altered fluid balance and evaporative water regulation. Despite comparable cumulative food intake between groups, DOCA-salt mice demonstrated significantly reduced locomotor activity, as measured by cumulative pedometer counts (**Figure 4D**), indicating suppressed spontaneous movement. Body weight did not differ significantly between DOCA-salt and

**Figure 4.**
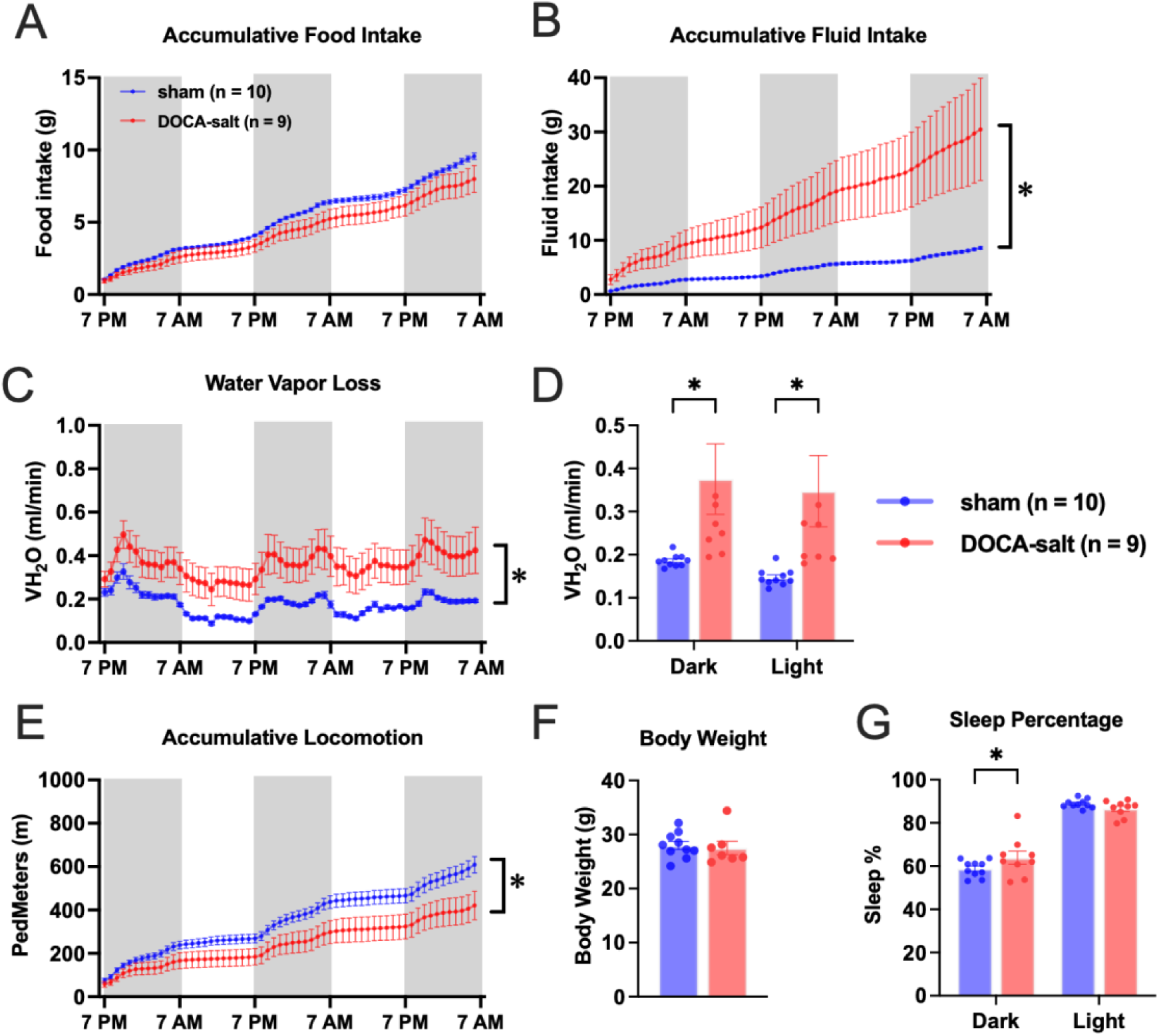
Effects of DOCA-salt treatment on metabolic and behavioral parameters. Mice were administered DOCA-salt (red; *n* = 9) or sham treated (blue; *n* = 10), and housed in metabolic cages for continuous monitoring from day 12 to 14 of DOCA or sham treatment. (A) Cumulative food intake. (B) Cumulative fluid intake (sham, water intake; DOCA-salt, 0.9% saline intake). (C, D) VH₂O loss, a proxy for thermoregulatory and respiratory water loss, over a 72-hour period (C) and average water vapor loss for 12/12 hours of dark and light cycles (D). (E) Locomotor activity measured as cumulative distance traveled (ped meters). (F) Body weight. (G) Average sleep percentage for 12/12 hours of dark and light cycles. Data are presented as means ± SEM (*P* < 0.05 vs. sham).

SHAM groups (**Figure 4E**). An analysis of sleep behavior revealed a modest, but significant, increase in total sleep percentage in DOCA-salt–treated mice (**Figure 4F**), consistent with decreased locomotor activity or increased rest during hypertension. There was no difference in energy expenditure, oxygen consumption (VO^2^), or carbon dioxide production (VCO^2^) between SHAM and DOCA-salt mice (**Figure 5A–I**). Notably, the respiratory exchange ratio (RER), an indicator of fuel utilization, was significantly lower in the DOCA-salt group during both light and dark phases (**Figure 5J, K**), suggesting a shift from carbohydrate metabolism toward lipid oxidation. Together, these findings demonstrate that DOCA-salt treatment affects water homeostasis, physical activity, sleep behavior, and fuel utilization, even in the absence of changes in caloric intake or body weight.

**Figure 5.**
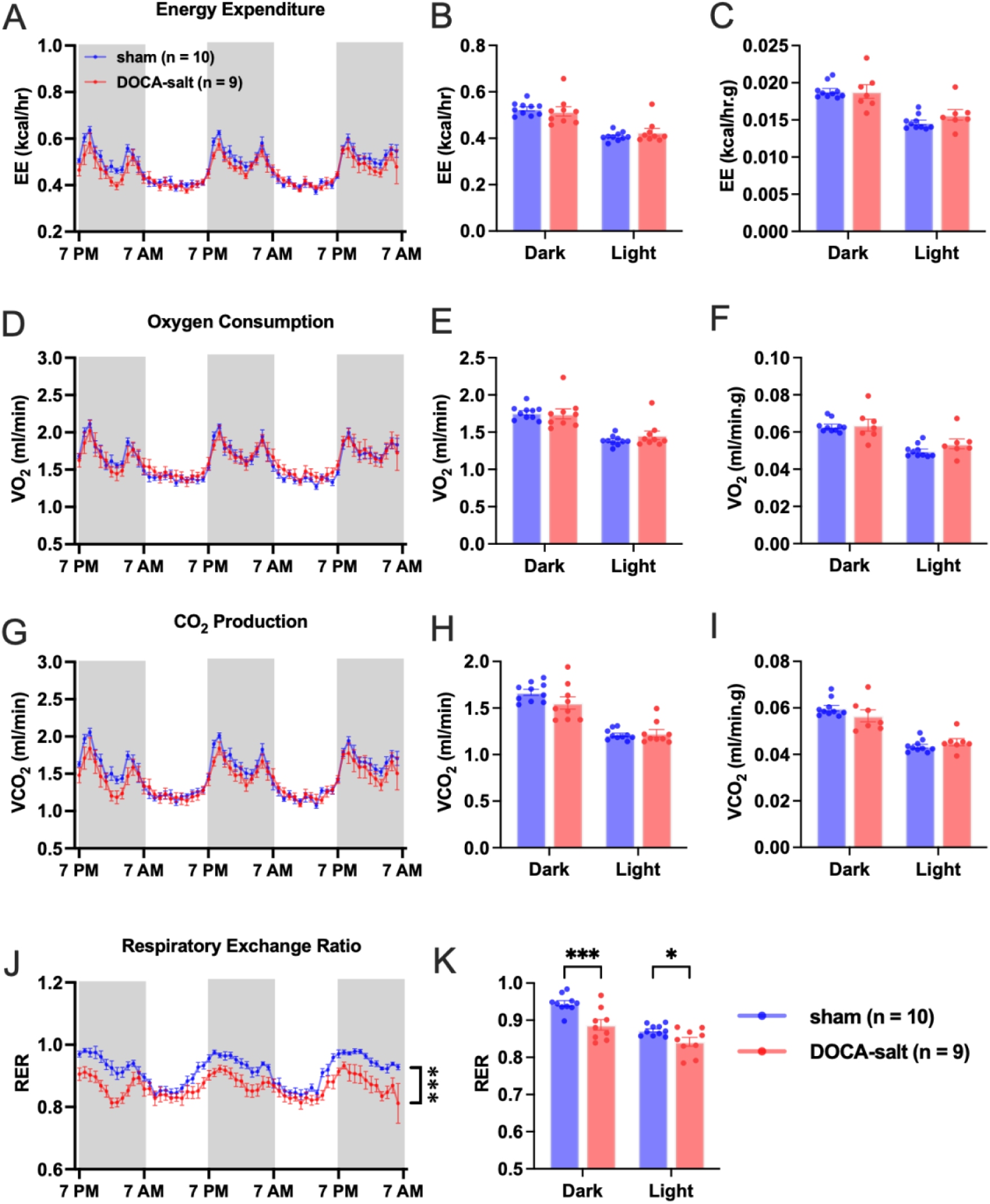
Effects of DOCA-salt treatment on whole-body energy metabolism and substrate utilization. Energy expenditure (EE), oxygen consumption (VO₂), carbon dioxide production (VCO₂), and respiratory exchange ratio (RER), measured in mice administered DOCA-salt (red; *n* = 9) or sham treated (blue; *n* = 10), and housed in metabolic cages for continuous monitoring from day 12 to 14 of DOCA or sham treatment. (A, B) Absolute EE over 72 hours (A) and an average over 12/12 hours of dark and light cycles (B). (C) EE normalized to body weight. (D, E) Absolute VO₂ over 72 hours (D) and an average over 12/12 hours of dark and light cycles (E). (F) weight-normalized VO₂. (G, H) VCO₂ levels over 72 hours (F) and an average over 12/12 hours of dark and light cycles (H). (I) VCO₂ normalized to body weight. (J, K) RER over 72 hours (J) and an average over 12/12 hours of dark and light cycles (K). Data are presented as means ± SEM (\**P* < 0.05, \*\*\**P* < 0.001 vs. SHAM).

### Cell Type-Specific Changes in the PVN of Mice Following DOCA-Salt Treatment

Next, we examined the impact of DOCA-salt treatment in PVN cells, performing an snRNA-seq analysis of dissociated PVN cells from SHAM or DOCA-salt mice (n = 10 males/group) after a 2-week treatment. A total of 4,908 cells from SHAM mice and 4,163 cells from DOCA-salt mice were sequenced. Following the first phase of the cell-type annotation pipeline in the Data Analysis section, we identified seven distinct clusters corresponding to major cell types—neurons, astrocytes, microglia, oligodendrocytes, endothelial cells and precursor oligodendrocytes—and a cluster of “other” cell populations (**Figure 6A**). Compared with SHAM mice, DOCA-salt–treated mice exhibited a significant shift in the percentage of oligodendrocytes (from 27.63% to 32.79%; *P* < 0.001) and neurons (from 55.7% to 51.0%; *P* < 0.001) (**Figure 6B**). A total of 2,734 neurons from SHAM mice and 2,121 neurons from DOCA-salt mice were sequenced, revealing eight distinct clusters of neuronal subtypes: TH neurons, cholinergic neurons, CRH neurons, GABAergic neurons, glutamatergic neurons, glutamatergic + GABAergic neurons, TRH neurons, and vasopressin neurons (**Figure 6C**). The percentage of GABAergic neurons in controls increased notably from 52.6% to 59.0% following DOCA-salt treatment (*P* < 0.001), whereas glutamatergic neurons decreased significantly from 35.2% to 23.2% (*P* < 0.001). The vasopressinergic neuronal subpopulation also significantly increased under DOCA-salt conditions to 12.0% (*P* < 0.001) compared with SHAM (**Figure 6D**). The observed decrease in glutamatergic neurons and the relative increase in GABAergic neurons in DOCA-salt hypertensive mice may reflect a compensatory neuroplastic response to chronic hypertension^38^, consistent with previous reports^39^. These findings demonstrate an increase in vasopressin neurons in response to DOCA-salt treatment, affirming that vasopressin neurons might be a key contributor to elevated blood pressure in DOCA-salt–treated animals.

**Figure 6.**
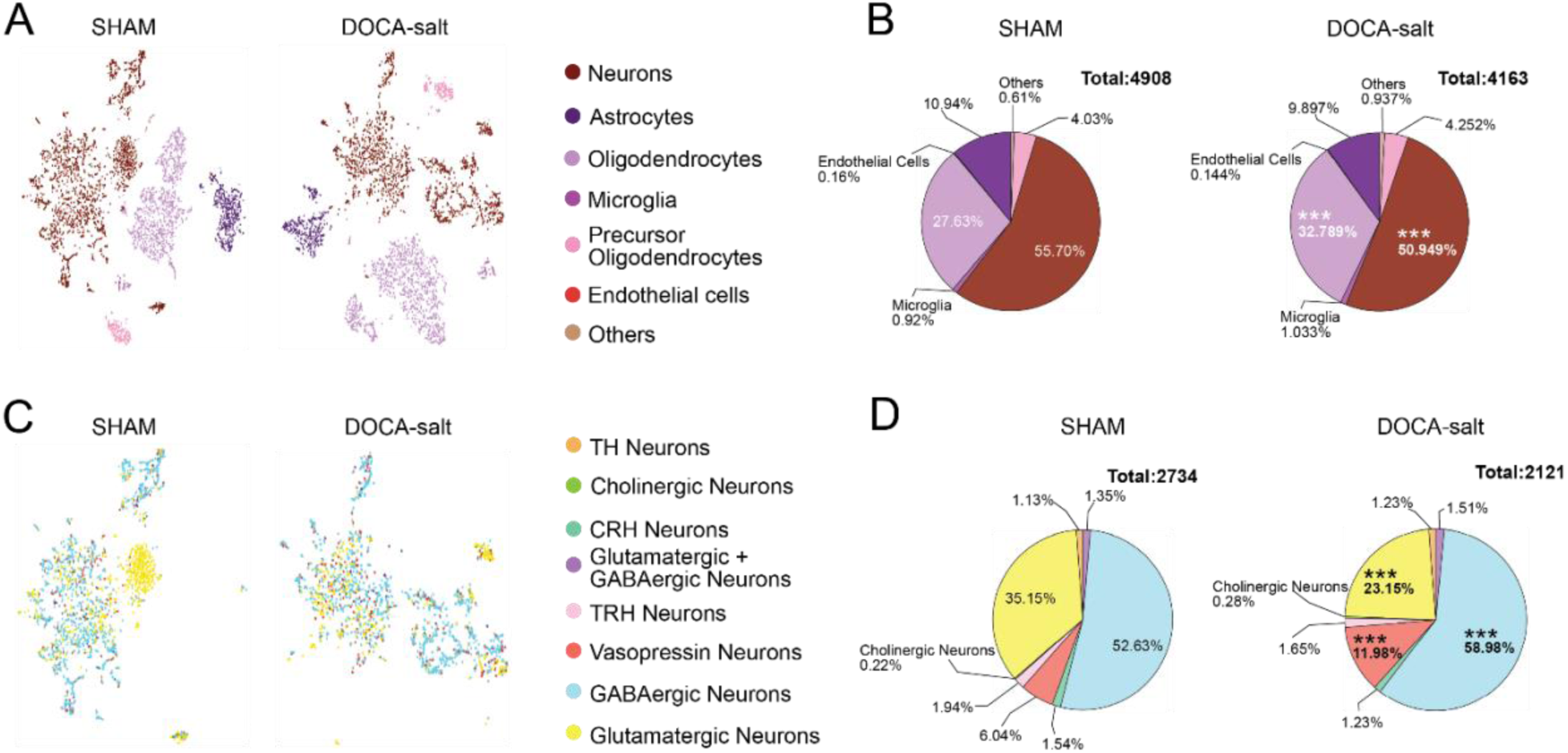
Impact of DOCA-salt treatment on cellular composition and neuronal subtypes in the PVN as revealed by snRNA-seq. (A) tSNE visualizations illustrating cell clustering and distribution of major PVN cell types. (B) Pie charts showing quantitative shifts in major cell populations due to DOCA-salt treatment, with significant increases in oligodendrocytes and decreases in neuronal proportions compared with SHAM. (C) tSNE visualizations depicting neuronal subtype distributions in SHAM and DOCA-salt treated mice. (D) Pie charts demonstrating significant alterations in neuronal subtype proportions under DOCA-salt conditions, with increases in GABAergic and vasopressin neurons and a marked reduction in glutamatergic neurons compared with SHAM. Colors correspond to neuronal subtypes and cell types, as indicated in the respective legends. Data are presented as means ± SEM (\*\*\**P* < 0.001 vs. SHAM; Fisher’s exact test).

### DOCA-Salt Enhances the Neuronal Vasoconstrictive RAS Axis While Reducing Astrocytic *Agt* in the PVN

To compare RAS gene expression between SHAM and DOCA-salt mice, we quantified the proportion of cell types expressing RAS genes, specifically determining the number of cells within each cell type that expressed a given RAS gene. For each RAS gene, the percentage of expressing cell types was calculated by dividing the number of cells of a particular type expressing the gene by the total number of cells expressing that RAS gene across the dataset. DOCA-salt treatment significantly decreased the percentage of *Agt-* expressing astrocytes (from 88.5% to 81.3%; *P* < 0.01) but increased the percentage of neurons expressing *Agt* (from 8.0% to 13.7%) and *Atp6ap*2 (from 77.0% to 80.7%; *P* < 0.01 for both) (**Figure 7**). No significant changes in expression of other RAS genes were observed across cell types. These findings provide evidence that DOCA-salt treatment activates the vasoconstrictive axis of RAS in neurons.

**Figure 7.**
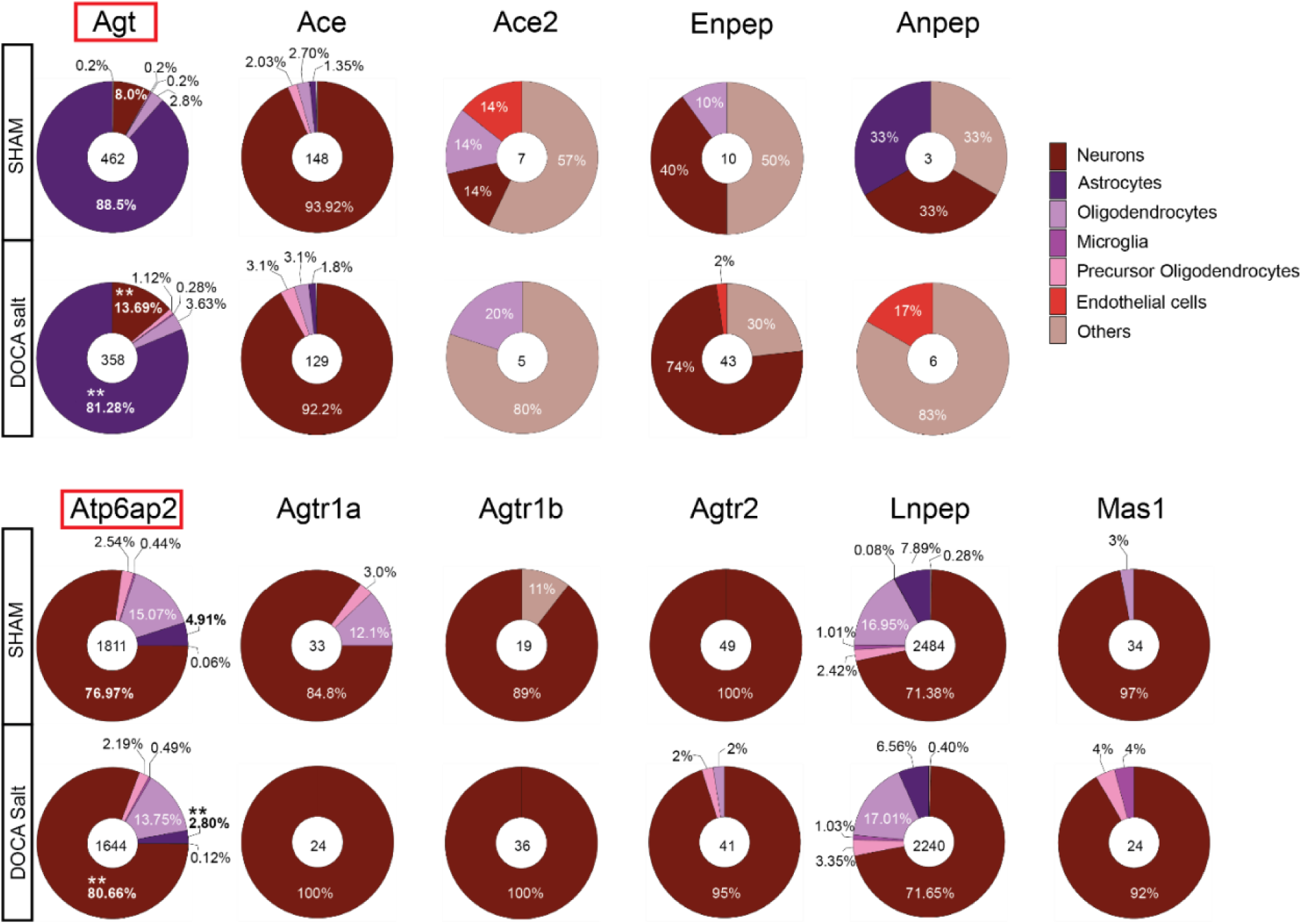
DOCA-salt treatment alters the cellular distribution of RAS gene expression in the PVN. Pie charts illustrating changes in the proportions of different PVN cell types expressing specific RAS genes in SHAM and DOCA-salt–treated mice. Numbers in the center of each pie chart represent the total counts of cells expressing each gene. Percentages indicate the proportion of each cell type among the cells expressing a given gene. Top panel: Shift in the cellular distribution of *Agt* expression with DOCA-salt treatment, with decreased proportions in astrocytes and increased proportions in neurons. Bottom panel: Shift in *Atp6ap2* expression following DOCA-salt treatment, with increased proportions of neurons expressing *Atp6ap2*. Colors denote cell types according to the provided legend. Data are presented as means ± SEM (\**P* < 0.01, \*\**P* < 0.001 for SHAM vs. DOCA-salt; Fisher’s exact test).

Among neuronal subpopulations, DOCA-salt treatment significantly increased the percentage of *Atp6ap2*-expressing vasopressin neurons (from 7.0% to 12.4%; *P* < 0.05) and decreased the percentage of *Atp6ap2-*expressing CRH neurons (from 1.9% to 1.0%; *P* < 0.01) and *Lnpep-*expressing glutamatergic neurons (from 22.4% to 18.9%; *P* < 0.05,) (**Figure 8**). The increase in *Atp6ap2*-expressing vasopressin neurons could be a key factor driving the hypertensive phenotype.

**Figure 8.**
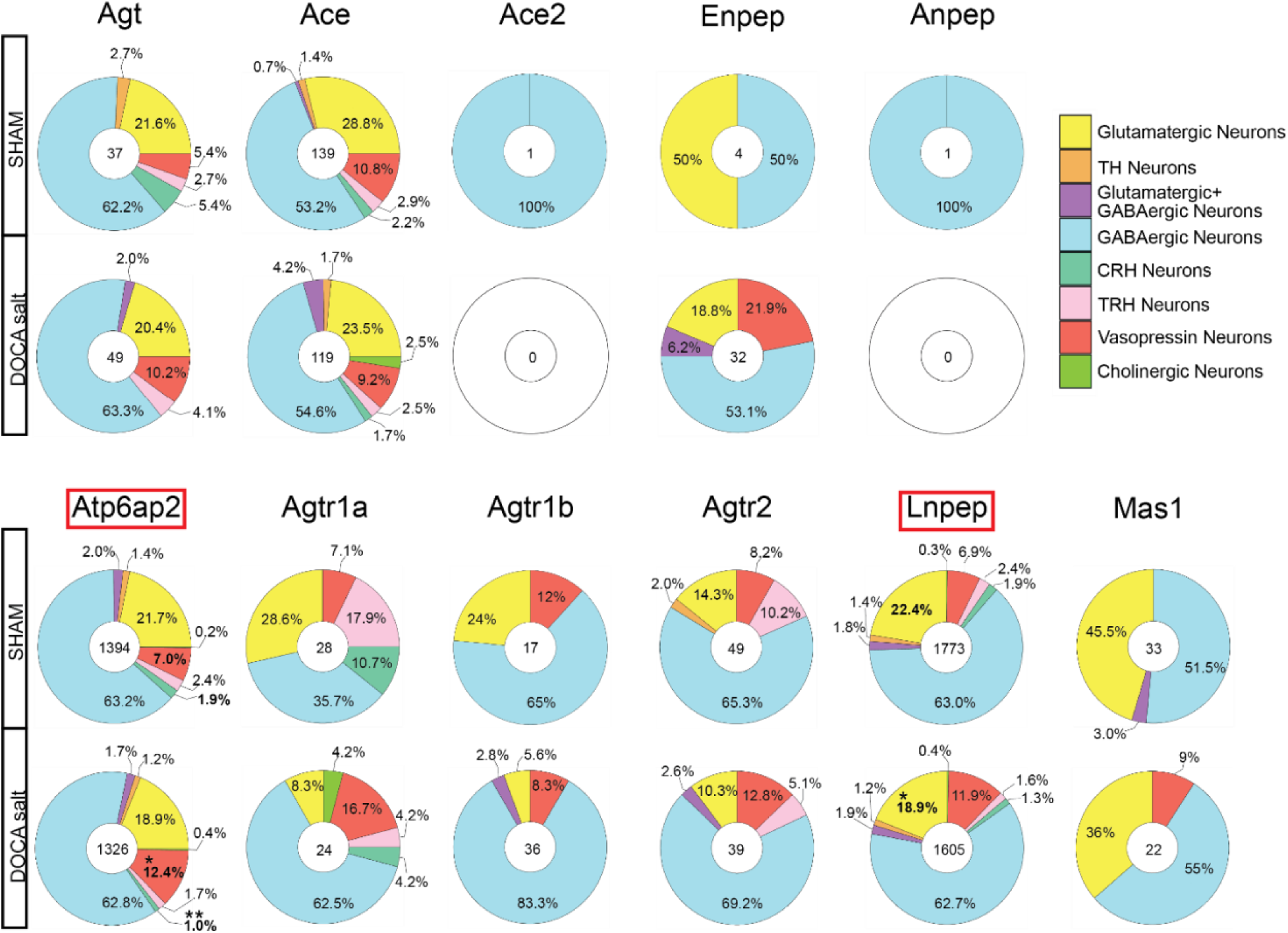
DOCA-salt treatment alters neuronal subtype-specific expression patterns of RAS genes in the PVN. Pie charts illustrating the proportions of neuronal subtypes expressing key RAS genes in SHAM and DOCA-salt–treated mice. The numbers in the center of each pie chart indicate the total number of neurons expressing each specific RAS gene, while percentages represent the distribution of neuronal subtypes among those cells. Top panel: Distribution of neuronal subtypes expressing the key substrate, AGT, and key enzymes for the RAS showing no significant subtype-specific alterations after DOCA-salt treatment. Bottom panel: Significant alterations in the subtype distribution of neurons expressing *Atp6ap2* under DOCA-salt conditions, with an increased proportion of vasopressin neurons and decreased proportions of CRH neurons and *Lnpep* (AT^4^R)-expressing glutamatergic neurons. Colors represent neuronal subtypes according to the provided legend. Data are presented as means ± SEM (\**P* < 0.05, \*\**P* < 0.01 vs. SHAM; Fisher’s exact test).

### Disrupted Energy Homeostasis and Behavioral Regulation Following HFD Feeding

To examine the metabolic and behavioral effects of HFD, at the end of the 6-week diet regimen, we housed mice in metabolic cages and continuously monitored them for 72 hours. Compared to CD-fed mice (n = 5), mice fed a HFD (n = 5) exhibited a significant increase in cumulative food intake (**Figure 9A**), consistent with hyperphagia associated with HFD feeding. HFD-fed mice showed significantly reduced water intake across the monitoring period (**Figure 9B**). In addition, VH₂O loss, a surrogate marker of evaporative water loss, was significantly reduced in the HFD group (**Figure 9C, D**), suggesting altered thermoregulatory or respiratory water handling. Despite increased caloric intake, HFD-fed mice demonstrated a marked reduction in locomotor activity, as evidenced by decreased cumulative distance traveled (**Figure 9E**). As expected, HFD-fed mice exhibited significantly higher body weight compared to CD-fed controls (**Figure 9F**). An analysis of sleep patterns revealed a significant increase in total sleep percentage during both dark and light phases in HFD-fed animals (**Figure 9G**), indicating altered behavioral rhythms and potential disruption of sleep-wake homeostasis.

**Figure 9.**
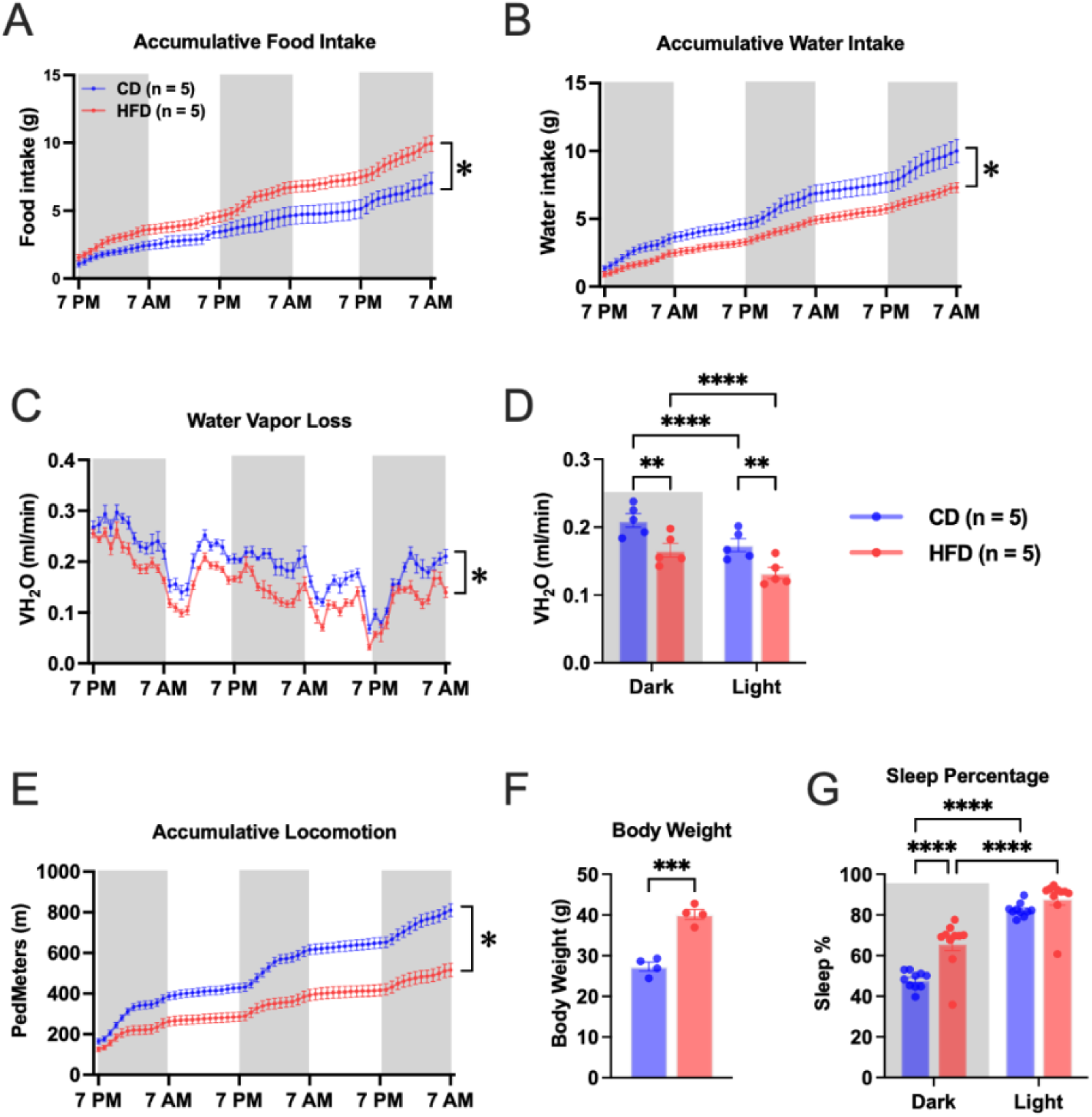
HFD alters metabolic and behavioral parameters. Metabolic analysis, performed over a 72-hour period at the end of a 6-week diet regimen in mice fed a HFD (60% calories from fat; n = 5) or CD (n = 5). (A) Cumulative food intake. (B) Cumulative water intake. (C, D) VH₂O loss, a proxy for thermoregulatory and respiratory water loss, over a 72-hour period (C) and average water vapor loss for 12/12 hours of dark and light cycles (D). (E) Locomotor activity measured as cumulative distance traveled (ped meters). (F) Body weight. (G) Average sleep percentage over 12/12 hours of dark and light cycles. Data are presented as means ± SEM (\*\**P* < 0.01, \*\*\**P* < 0.001, \*\*\*\**P* < 0.0001 vs. CD).

HFD feeding resulted in a small, but significant, increase in absolute energy expenditure (EE) across both dark and light phases (**Figure 10A, B**). Normalized to body weight, EE was significantly lower in HFD-fed mice (**Figure 10C**), suggesting diet-induced suppression of metabolic rate. Absolute oxygen consumption (VO₂) was increased in HFD-fed mice across the circadian cycle relative to those in CD-fed controls (**Figure 10D- E**). However, normalized to body weight, VO₂ was significantly lower in HFD-fed mice (**Figure 10F**). Carbon dioxide production (VCO₂) was significantly lower in the HFD group whether expressed as absolute or normalized values (**Figure 10G–I**). Notably, RER, an indicator of substrate utilization, was significantly lower in HFD-fed mice during both dark and light phases (**Figure 10J, K**), a reduction that reflects a metabolic shift toward increased lipid oxidation and reduced carbohydrate utilization in response to HFD feeding.

**Figure 10.**
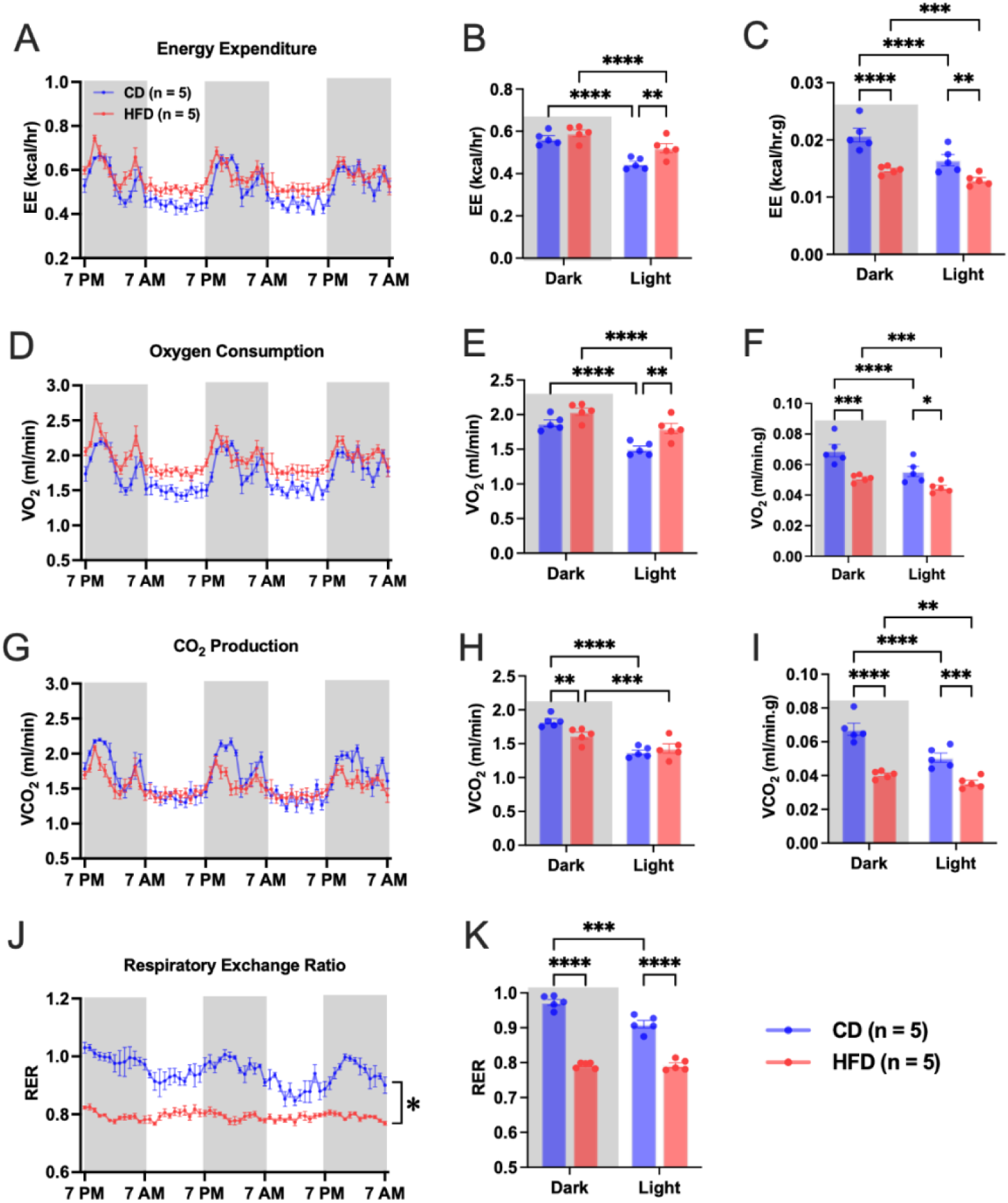
HFD alters whole-body energy metabolism and substrate utilization. Energy expenditure (EE), oxygen consumption (VO₂), carbon dioxide production (VCO₂), and respiratory exchange ratio (RER) measured in mice fed a CD or HFD for 6 weeks. (A, B) Absolute EE over 72 hours (A) and an average over 12/12 hours of dark and light cycles (B). (C) EE normalized to body weight. (D, E) Absolute VO₂ over 72 hours (D) and an average over 12/12 hours of dark and light cycles (E). (F) weight-normalized VO₂. (G, H) VCO₂ levels over 72 hours (G) and an average over 12/12 hours of dark and light cycles (H). (I) VCO₂ normalized to body weight. (J, K) RER over 72 hours (J) and an average over 12/12 hours of dark and light cycles (K). Data are presented as means ± SEM (\**P* < 0.05, \*\**P* < 0.01, \*\*\**P* < 0.001, \*\*\*\**P* < 0.0001 vs. CD).

Collectively, these findings demonstrate that short-term HFD feeding leads to coordinated alterations in energy intake, fluid consumption, physical activity, evaporative water loss, and sleep behavior. In parallel, HFD feeding suppresses energy expenditure and induces a metabolic shift toward increased fat oxidation.

### HFD Increases Excitatory and Stress-response Signals While Reducing Inhibitory Signals in the PVN

To understand the impact of HFD feeding on PVN cellular phenotypes, we performed snRNA-seq on isolated PVN single nuclei from mice fed a CD or HFD (n = 10 males/group) for 6 weeks. A total of 3,652 cells from CD mice and 4,477 cells from HFD mice were sequenced. A cells annotation pipeline performed as in previous experiments (above) identified seven distinct clusters corresponding to the same seven major cell types (**Figure 11A**). Compared with CD mice, HFD-fed mice exhibited a significant shift in the percentage of oligodendrocytes (from 34.5% to 31.9%; *P* < 0.05) and neurons (from 44.8% to 49.1%; *P* < 0.001) (**Figure 11B**). A total of 1,637 neuronal subpopulations from CD mice and 2,200 neuronal subpopulations from HFD mice were sequenced, again revealing the same eight distinct neuronal subtypes (**Figure 11C**). HFD feeding significantly altered the percentage of specific neuronal subtypes, reducing GABAergic neurons (from 54.1% to 46.7%; *P* < 0.001), while increasing glutamatergic neurons (from 27.7% to 30.9%; *P* < 0.05), vasopressin neurons (from 9.5% to 13.0%; *P* < 0.05), and CRH neurons (from 1.0% to 2.6%; *P* < 0.05) (**Figure 11D**). These data suggest that HFD treatment shifts the balance toward increased excitatory and stress-responsive signaling, while reducing inhibitory signals in the PVN.

**Figure 11.**
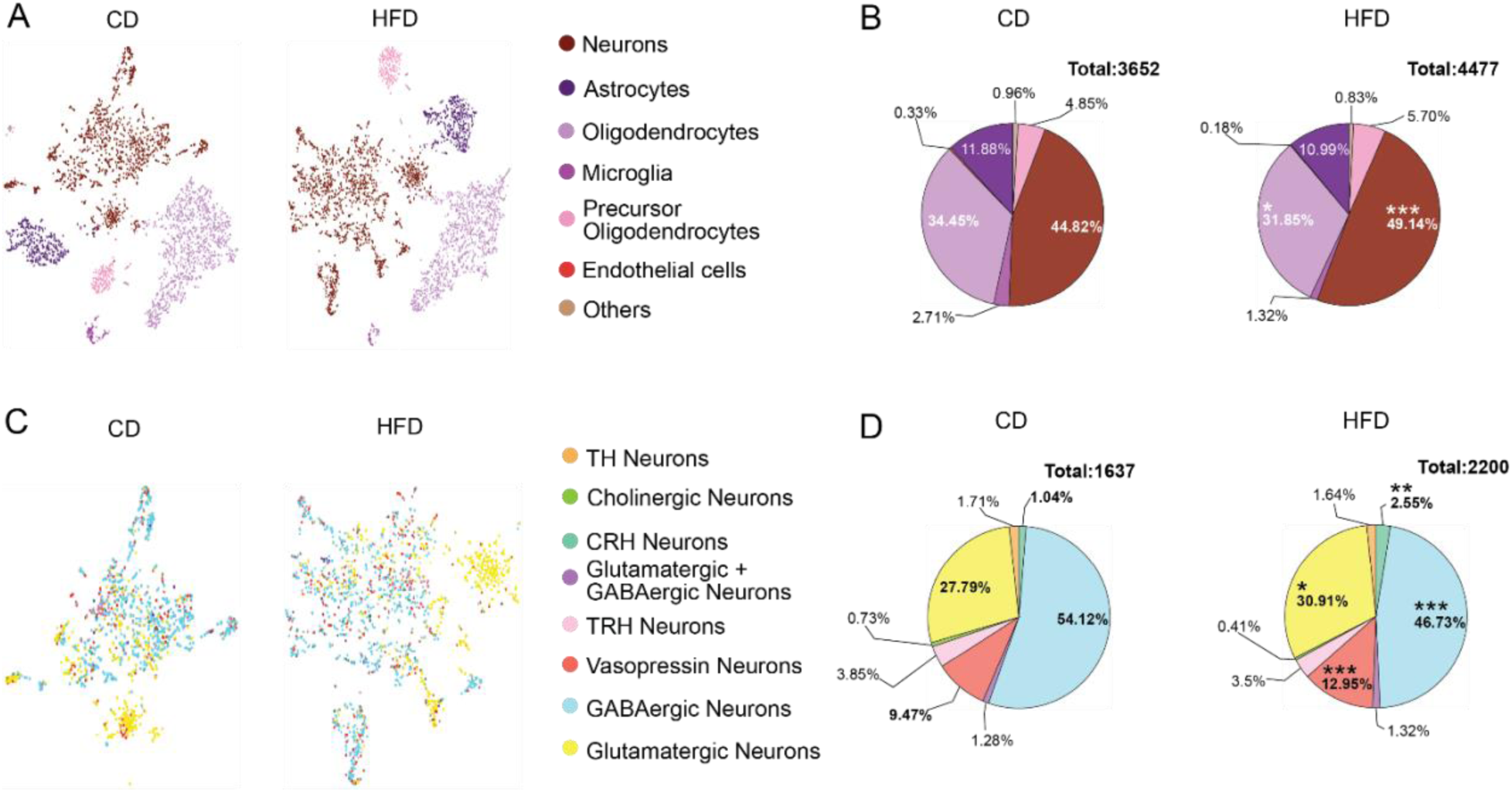
Effects of a HFD on the cellular composition and neuronal subtype distribution in the PVN revealed by snRNA-seq. (A) tSNE visualizations displaying the clustering of major PVN cell types in CD- and HFD-fed mice. (B) Pie charts indicating significant quantitative shifts in major cell populations under HFD conditions, with significantly decreased proportions of oligodendrocytes and increased proportions of neurons. (C) tSNE visualizations illustrating neuronal subtype distributions between CD and HFD mice. (D) Pie charts demonstrating significant changes in neuronal subtype proportions under HFD, including decreases in GABAergic neurons and increases in glutamatergic neurons and vasopressin neurons. Colors denote cell and neuronal subtypes according to the provided legend. Data are presented as means ± SEM (\**P* < 0.05, \*\**P* < 0.01, \*\*\**P* < 0.001 for CD vs. HFD; Fisher’s exact test).

### HFD Induces Cell-Type-Specific Alterations in Brain RAS Gene Expression

To compare the cellular distribution of RAS-related genes between CD- and HFD-fed mice, we calculated the percentage of cell types expressing RAS genes using the same method described above. HFD treatment increased the percentage of *Lnpep*-expressing neurons (from 62.8% to 66.2%; *P* < 0.05), but, interestingly, slightly decreased the percentage of microglia expressing *Atp6ap2* (from 2.5% to 0.8%) or *Lnpep* (from 3.4% to 1.2%; *P* < 0.001 for both) (**Figure 12**). The increase in *Lnpep*-expressing neurons and reduction in *Atp6ap2-* and *Lnpep-*expressing microglia likely reflects an alteration in cellular signals of the local RAS within the brain under HFD conditions.

**Figure 12.**
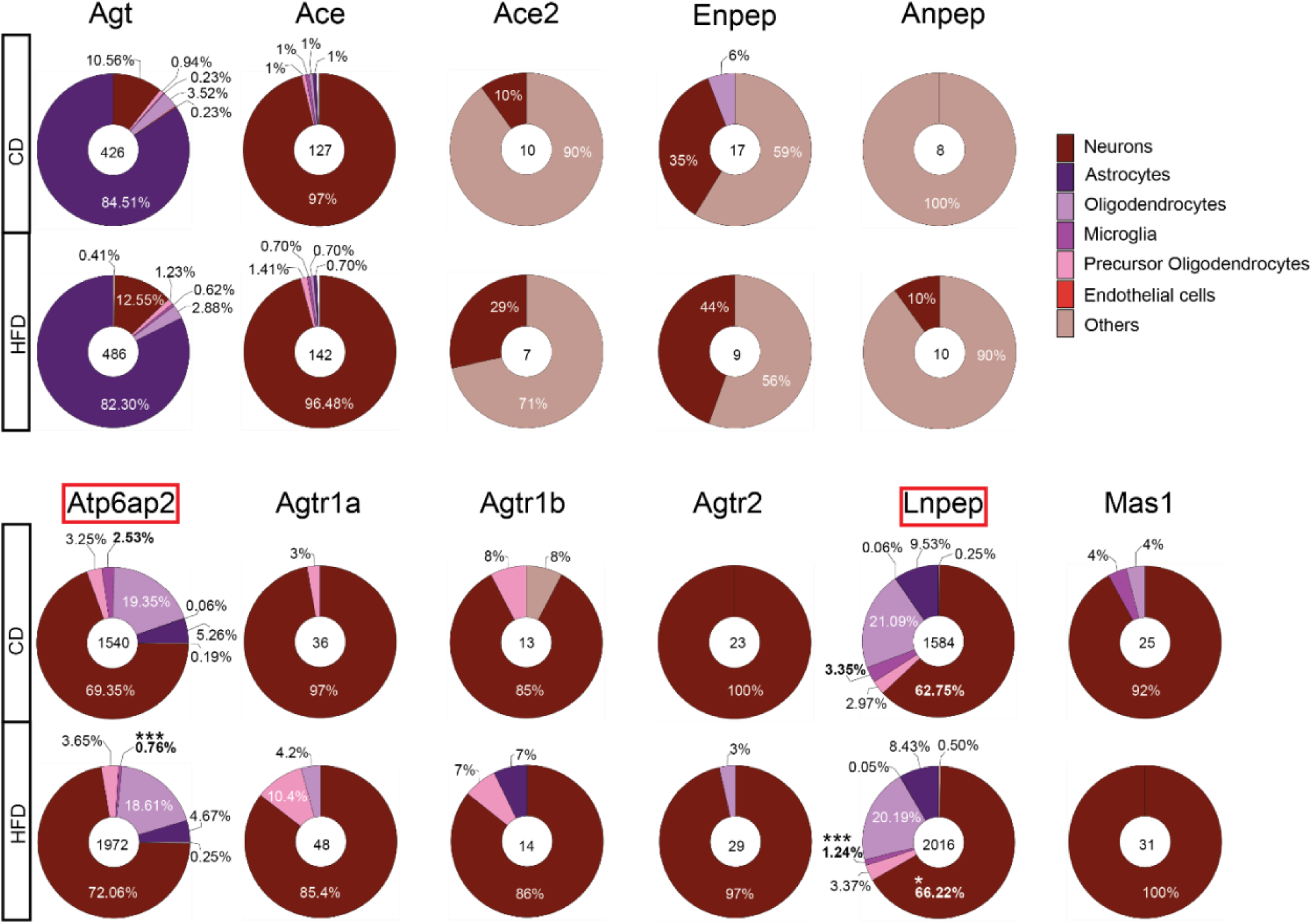
HFD induces cell-type-specific alterations in RAS gene expression in the PVN. Pie charts illustrating the distribution of PVN cell types expressing specific RAS genes in CD- versus HFD-fed mice. Numbers in the center of each pie chart indicate total counts of cells expressing each RAS gene, and percentages represent the proportion of each cell type among cells expressing a given gene. Top panel: Proportions of cell types expressing *Ag*t and genes encoding key RAS enzymes, showing no significant shifts under HFD conditions. Bottom panel: Significant increase in the proportion of *Lnpep*- expressing neurons under HFD, together with slight but significant reductions in *Atp6ap2*- and *Lnpep*-expressing microglia. Colors denote specific cell types, as indicated in the provided legend. Data are presented as means ± SEM (\**P* < 0.05, \*\*\**P* < 0.001 for CD vs. HFD; Fisher’s exact test).

A neuronal subpopulation analyses showed that a 6-week HFD regimen significantly decreased the percentage of *Agt-, Atp6ap2-*, *Agtr1b-, Lnpep-,* and *Mas1**-***expressing GABAergic neurons (*P* < 0.05) (**Figure 13**). Notably, HFD feeding significantly increased the percentage of vasopressin neurons expressing *Atp6ap2* (from 10.02% to 14.99%), *Agtr1b* (from 0% to 42.7%), *Lnpep* (from 9.3% to 14.5%) or *Mas1* (from 0% to 22.6%; *P* < 0.05 for each). The percentage of *Atp6ap2*-expressing CRH neurons was also slightly increased (from 1.7% to 3.1%; *P* < 0.05). These data indicate that HFD significantly impacts RAS-expressing neuronal subpopulations, especially GABAergic and vasopressin neurons.

**Figure 13.**
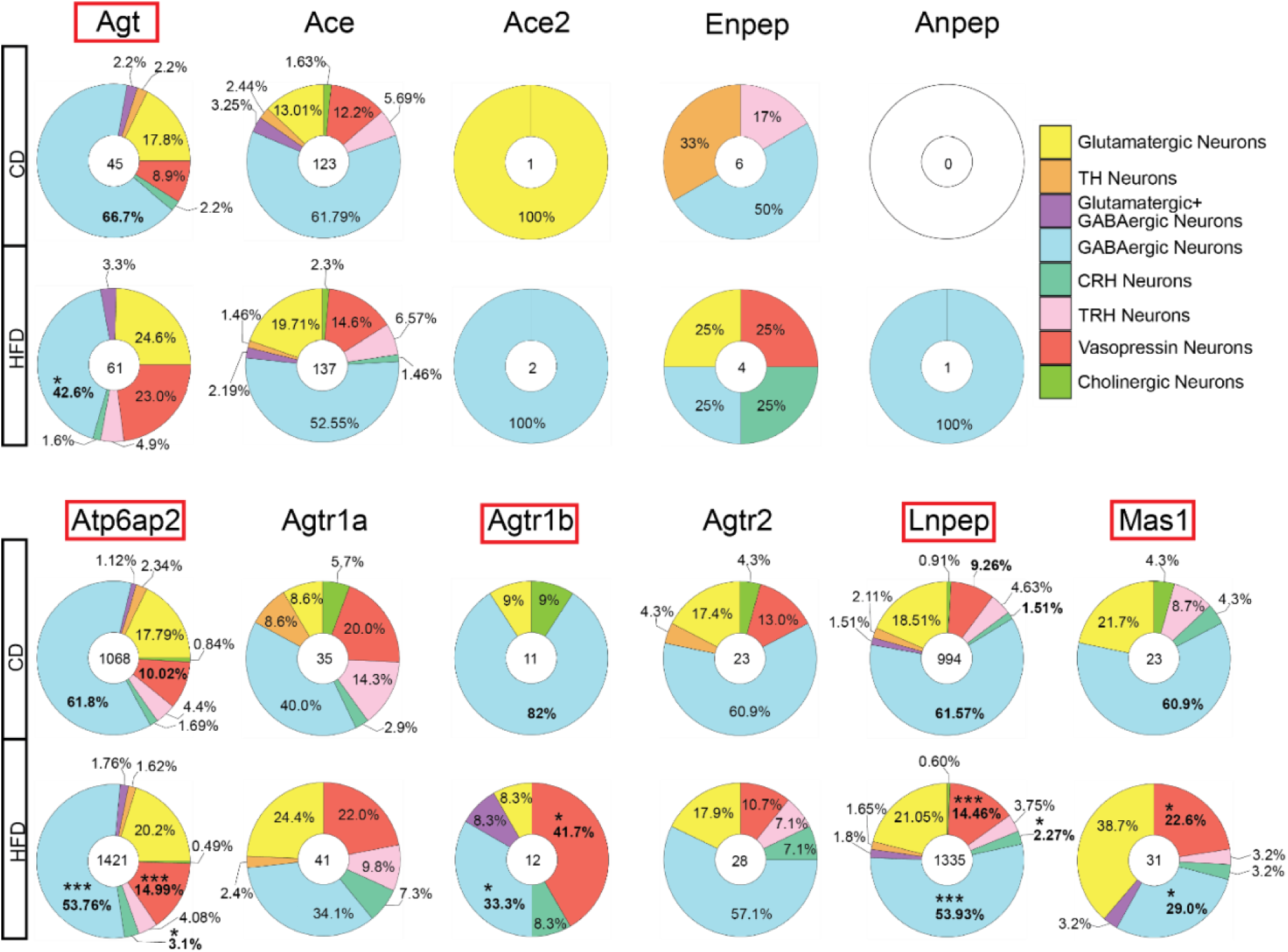
HFD induces neuronal subtype-specific alterations in RAS gene expression in the PVN. Pie charts illustrating the distribution of neuronal subtypes expressing specific RAS genes in CD- versus HFD-fed mice. Numbers in the center of each pie chart indicate the total counts of neurons expressing the respective RAS gene, and percentages represent the proportion of each neuronal subtype among neurons expressing each gene. Top panel: Subtype-specific expression patterns of *Agt* and other RAS genes, demonstrating notable reductions in the proportion of GABAergic neurons expressing *Agt* under HFD conditions. Bottom panel: Significant subtype-specific alterations under HFD, including decreased proportions of GABAergic neurons expressing *Atp6ap2*, *Agtr1b*, *Lnpep*, and *Mas1*, together with marked increases in vasopressin neuronal populations expressing these genes. Colors correspond to neuronal subtypes indicated in the provided legend. Data are presented as means ± SEM (\**P* < 0.05, \*\*\**P* < 0.001 for CD vs. HFD; Fisher’s exact test).

## Discussion

In this study, we used snRNA-seq to provide a comprehensive analysis of the cellular and transcriptomic landscape of the RAS in the PVN of male mice. By examining the effects of a DOCA-salt or HFD regimen, we elucidate critical cellular and molecular adaptations of the RAS in the PVN—a key brain region for blood pressure regulation and energy homeostasis. We confirm at the single-cell level that expression of RAS genes in the brain hypothalamus show strong cell-type specificity: *Agt* was enriched in astrocytes, while other key RAS-related genes, including *Ace*, *Atp6ap2*, *Agtr1a*, *Lnpep* and *Mas1,* were predominantly expressed in neurons. Neuronal subtypes, including GABAergic, glutamatergic, vasopressin and CRH neurons, displayed distinct RAS gene expression patterns. DOCA-salt treatment increased the proportion of GABAergic and vasopressin neurons; it also enhanced the percentage of *Atp6ap2-* and *Agt*-expressing neurons while reducing *Agt*-expressing astrocytes, suggesting activation of a vasoconstrictive RAS axis in hypertensive states. HFD increased the neuronal proportion and shifted the neuronal subtype distribution by enhancing excitatory (glutamatergic) and stress-responsive (vasopressin, CRH) neurons while reducing inhibitory (GABAergic) neurons. HFD also induced cell-type-specific changes in RAS gene expression, notably increasing the proportion of *Atp6ap2-*, *Agtr1b-*, *Lnpep-*, and *Mas1*-expressing vasopressin neurons while reducing GABAergic neurons expressing multiple RAS components. These changes suggest that both DOCA-salt and HFD treatments remodel RAS signaling in a cell-type- and subtype-specific manner, potentially contributing to neuroendocrine and autonomic dysfunction.

Our initial snRNA-seq analysis revealed distinct cell type-specific expression patterns of RAS genes within the PVN. Neurons exhibited the most diverse expression profiles, indicative of their central role in RAS signaling, showing expression of *Agt, Ace, Atp6ap2*, *Agtr1a*, *Agtr1b*, *Agtr2*, *Lnpep*, and *Mas1*. Astrocytes showed high *Agt* expression, consistent with previous findings^40–42^, supporting their function as a key source of local angiotensinogen and paracrine modulation within the brain. These results reinforce the role of the PVN as a critical hub for RAS activity that contributes to blood pressure regulation and neuroendocrine control^43–46^. Although all core RAS components have been reported in the brain^47^, the presence of *Agtr2* in the PVN has been debated, with some studies failing to detect it^48, 49^. Our findings, together with others^50, 51^, support *Agtr2* expression in the PVN, with discrepancies across studies likely reflecting differences in methodology, such as tissue processing or detection sensitivity. Further investigation is warranted to clarify the expression dynamics and functional significance of *Agtr2* in this region. Despite the recognized role of renin (*Ren*) in the brain RAS, we were unable to detect *Ren* mRNA in our snRNA-seq dataset. This may be attributable to several technical and biological factors. First, the expression of renin in the brain is known to be extremely low and highly localized^26, 52^, making it challenging to capture with snRNA-seq, which generally samples a limited number of nuclei per region and may miss rare transcripts. Second, nuclear RNA represents a subset of total cellular RNA, and low-abundance transcripts such as *Ren* may fall below the detection threshold, particularly if their nuclear localization is transient or minimal. Therefore, while our dataset provides important insights into RAS component expression at the single-nucleus level, the sensitivity of the underlying technique is insufficient to detect *Ren* expression in hypothalamic regions.

Neuronal subtype analysis identified eight distinct populations with unique RAS gene expression patterns, suggesting specialized stress-response and fluid-homeostasis roles. Notably, *Anpep*, encoding alanyl aminopeptidase, an enzyme involved in converting angiotensin III to angiotensin IV, was absent in all neuronal populations, consistent with prior transcriptomic and immunohistochemical studies^53, 54^. Given that Ang IV exerts hypertensive effects via LNPEP^55^, the lack of ANPEP in neurons suggests that Ang IV production may occur via non-neuronal sources or through alternative enzymatic pathways, potentially involving astrocytic or endothelial signaling.

DOCA-salt treatment induced notable changes in PVN cellular composition, characterized by an increased proportion of oligodendrocytes and a reduction in neurons. Within the neuronal population, there was a marked increase in GABAergic and vasopressin neurons, together with a significant decline in glutamatergic neurons. These findings are consistent with previous reports showing that DOCA-salt enhances GABAergic excitation in vasopressin neurons^56^ and increases vasopressin expression and signals in DOCA-salt hypertension^37, 57, 58^. The finding of an increase in the GABAergic neuronal population in the DOCA-salt hypertension model was interesting but unsurprising. There have been conflicting reports on the changes in the GABAergic system in the PVN of hypertensive animals depending on the hypertension stage, hypertension models, and techniques used (e.g., molecular vs. electrophysiology)^59, 60^. Li et al. reported increased tonic GABAergic inhibition of PVN parasympathetic neurons through the GABA^B^ receptor in 13-week-old spontaneously hypertensive rats (established stage of hypertension) but not in normotensive rats^61^, consistent with our finding of an increase in the GABAergic neuronal population in the established stage of hypertension (2-week DOCA-salt treatment). Interestingly, we observed a reduction in the percentage of glutamatergic neurons within neuronal subpopulations following 2 weeks of DOCA-salt treatment. Glutamatergic neurons in the PVN mediate an increase in blood pressure in DOCA-salt hypertension^62^. Thus, such shifts may represent adaptive or maladaptive responses that serve to maintain excitatory/inhibitory balance in the face of chronic hypertensive stress. Changes in the cellular expression pattern of RAS genes under DOCA-salt conditions further support this interpretation. The upregulation of *Agt* and *Atp6ap2* in neurons—especially in vasopressin neurons—suggests an enhanced RAS-driven hypertensive signaling pathway in vasopressin neurons^57, 63^. Conversely, the reduction in *Atp6ap2* expression in CRH neurons and *Lnpep* expression in glutamatergic neurons may reflect compensatory mechanisms aimed at dampening stress and excitatory responses.

HFD feeding similarly led to significant alterations in PVN cell composition, with increased neuronal and decreased oligodendrocyte proportions. Oligodendrocytes are essential for myelin production and efficient neuronal communication^64–66^, and their vulnerability to HFD-induced oxidative stress and mitochondrial dysfunction has been well documented^67, 68^. Loss of oligodendrocytes and disrupted myelination may impair neural network function and contribute to HFD-induced pathophysiology. At the neuronal subtype level, a HFD increased the proportions of glutamatergic, vasopressin and CRH neurons, suggesting a shift toward heightened excitatory and stress-responsive signaling. Enhanced glutamatergic excitability has been implicated in obesity-related neural dysregulation^69, 70^, while CRH neurons play a central role in initiating the hypothalamic-pituitary-adrenal (HPA) axis stress response. Although a HFD has been shown to blunt CRH activity, the observed increase in CRH neuron numbers could reflect a compensatory or dysregulated response^71^. Vasopressin neurons—critical for regulating fluid balance, stress and blood pressure^5, 72^—also expanded under HFD, though direct evidence linking a HFD to altered vasopressin neuron function remains limited and warrants further study.

Importantly, RAS gene expression analyses revealed cell-type-specific remodeling under a HFD. *Atp6ap2-*, *Agtr1b-*, *Lnpep-*, and *Mas1*-expressing vasopressin neurons were significantly elevated, pointing to their potential involvement in hypertension and metabolic dysfunction. In contrast, reduced expression of multiple RAS genes—including *Agt, Atp6ap2*, *Agtr1b*, *Lnpep*, and *Mas1*—in GABAergic neurons suggests a diminished inhibitory RAS component, potentially exacerbating PVN excitability and sympathetic activation.

Our metabolic cage data allowed us to uncover physiological functions that correlated with transcriptional changes observed in the PVN. DOCA-salt–treated mice exhibited a significant increase in water intake and water vapor loss, together with reduced locomotor activity and increased sleep duration, despite unchanged caloric intake and body weight, suggesting a phenotype of behavioral slowing and altered fluid balance. In contrast to a previous report^73^, we found no change in energy expenditure or oxygen consumption. However, the respiratory exchange ratio was significantly reduced, indicating a metabolic shift favoring lipid utilization. The discrepancy in energy expenditure findings may be attributable to differences in the timing of data collection, as our measurements were obtained 12–14 days after DOCA-salt treatment, whereas the previous study assessed animals at 21 days post treatment^73^. The metabolic phenotype of the DOCA-salt model aligns with the increased proportion of vasopressin neurons and altered RAS gene expression patterns, including elevated *Atp6ap2* expression in vasopressin neurons. Together, these results support previous findings that DOCA-salt disrupts fluid regulation and energy utilization in a manner linked to PVN vasopressinergic remodeling^57, 74^ and activation of the vasoconstrictive arm of the brain RAS^13, 75, 76^. The incorporation of metabolic cage phenotyping extends the significance of our snRNA-seq findings by establishing functional readouts for the observed cellular and transcriptional plasticity. These data demonstrate that RAS remodeling in the PVN is correlated with behavioral and metabolic outputs, underscoring the role of the PVN as a neuroendocrine interface through which dietary and hormonal stressors drive maladaptive autonomic and metabolic responses.

In contrast, HFD-fed mice exhibited a distinct metabolic phenotype characterized by hyperphagia, reduced water intake, lower evaporative water loss, and decreased locomotion. HFD mice also displayed increased body weight and total sleep percentage, features consistent with disrupted behavioral rhythms and reduced energy expenditure^77, 78^. Notably, while absolute energy expenditure and VO₂ were slightly elevated in HFD-fed mice, normalization of these values to body weight revealed suppressed metabolic efficiency and significantly reduced RER values, pointing to an HFD-induced shift toward fat oxidation. These physiological responses were correlated with PVN transcriptomic changes, including increased proportions of excitatory (glutamatergic) and stress-related (CRH, vasopressin) neurons, and decreased proportions of inhibitory (GABAergic) neurons.

This study provides an extensive characterization of cellular composition and RAS gene expression patterns in the PVN. However, it is not without limitations. First, fresh tissue dissections used for sequencing included the PVN region but could have encompassed periventricular and surrounding hypothalamic areas. Consequently, the obtained transcriptional profiles may not exclusively represent PVN-specific populations, potentially influencing the interpretation of cell-type-specific changes in RAS gene expression. Secondly, although the study included a relatively large number of animals (n = 10/group), only ∼5000 cells per experimental condition were sequenced. While this provides valuable insights, a higher sequencing depth and increased cell count could improve resolution, enable the detection of less abundant cell types and enhance statistical power, potentially revealing additional subtle but biologically relevant differences. Third, the current analyses were exclusively performed using male mice. Given known sex differences in cardiovascular and metabolic regulation^79, 80^ as well as neuroendocrine function, these results may not fully represent the transcriptomic landscape or RAS gene dynamics in females. Future studies including both sexes would provide critical insights into sex-dependent mechanisms in RAS-mediated cardiovascular and metabolic regulation. Lastly, while the transcriptomic data provide comprehensive information regarding cell-type-specific changes and neuronal subtype-specific RAS gene expression, functional validation is still required. Further physiological and pharmacological experiments should be conducted to confirm the functional roles of these identified cell populations and RAS components in regulating blood pressure, metabolism, and neuroendocrine responses. Addressing these limitations in future research will further clarify the role of cell-type-specific RAS signaling in the PVN and enhance the translational potential of these findings in clinical contexts.

Additionally, the current study employed single-nucleus rather than single-cell RNA-seq. While snRNA-seq offers advantages in capturing transcriptional profiles from frozen or difficult-to-dissociate tissues and reduces transcriptional artifacts introduced during enzymatic dissociation, it primarily captures nuclear rather than cytoplasmic RNA. Consequently, snRNA-seq may underrepresent certain transcripts, particularly those localized predominantly in the cytoplasm, potentially missing key biological signals detectable by traditional scRNA-seq methods. Future comparisons using both approaches could further enhance the comprehensive understanding of cellular transcriptional dynamics in the PVN.

Together, these findings highlight the dynamic interplay among RAS signaling, cellular composition, and disease state within the PVN. Both DOCA-salt and HFD trigger distinct yet overlapping patterns of neuronal remodeling and RAS gene expression, aligning with their roles in driving neuroendocrine and cardiovascular dysfunction. This study underscores the utility of single-cell transcriptomics in dissecting the molecular underpinnings of neurogenic hypertension and obesity. Future investigations should aim to functionally characterize specific neuronal and glial subtypes to better understand their contributions to RAS-mediated pathophysiology and explore novel cell-targeted therapies for cardiometabolic diseases.

## Acknowledgements

We thank Silvana Cooper for the technical support on the project.

## Funding

This work was supported, in part, by grants from the National Institutes of Health (NIH; R01HL122770, R01DK135621, R35HL122770) to Y. Feng Earley, and from the National Science Foundation (NSF; 343019, 2203236), National Institute of General Medical Sciences (NIGMS; R44GM152152), National Cancer Institute (NCI; U01CA274573), and National Institute of Food and Agriculture (NIFA; 2023-6702240041) to T. Nguyen.

## Notes

### Competing Interest Statement

The authors have declared no competing interest.

## References

1. Benarroch EE. Paraventricular nucleus, stress response, and cardiovascular disease. Clin Auton Res. 2005;15 (4):254–263. doi: 10.1007/s10286-005-0290-7.

2. Pan S, Worker CJ, Feng Earley Y. The hypothalamus as a key regulator of glucose homeostasis: emerging roles of the brain renin-angiotensin system. Am J Physiol Cell Physiol. 2023;325 (1):C141–C154. doi: 10.1152/ajpcell.00533.2022.

3. Carmichael CY, Wainford RD. Hypothalamic signaling mechanisms in hypertension. Curr Hypertens Rep. 2015;17 (5):39. doi: 10.1007/s11906-015-0550-4.

4. Ferguson AV, Latchford KJ, Samson WK. The paraventricular nucleus of the hypothalamus - a potential target for integrative treatment of autonomic dysfunction. Expert Opin Ther Targets. 2008;12 (6):717–727. doi: 10.1517/14728222.12.6.717.

5. Savic B, Murphy D, Japundzic-Zigon N. The Paraventricular Nucleus of the Hypothalamus in Control of Blood Pressure and Blood Pressure Variability. Front Physiol. 2022;13:858941. doi: 10.3389/fphys.2022.858941.

6. Khor S, Cai D. Hypothalamic and inflammatory basis of hypertension. Clin Sci (Lond). 2017;131 (3):211–223. doi: 10.1042/CS20160001.

7. Herrera Moro Chao D, Kirchner MK, Pham C, et al. Hypothalamic astrocytes control systemic glucose metabolism and energy balance. Cell Metab. 2022;34 (10):1532–1547 e1536. doi: 10.1016/j.cmet.2022.09.002.

8. Xi H, Li X, Zhou Y, Sun Y. The Regulatory Effect of the Paraventricular Nucleus on Hypertension. Neuroendocrinology. 2024;114 (1):1–13. doi: 10.1159/000533691.

9. Chen F, Cham JL, Badoer E. High-fat feeding alters the cardiovascular role of the hypothalamic paraventricular nucleus. Am J Physiol Regul Integr Comp Physiol. 2010;298 (3):R799–807. doi: 10.1152/ajpregu.00558.2009.

10. Zhang ZH, Francis J, Weiss RM, Felder RB. The renin-angiotensin-aldosterone system excites hypothalamic paraventricular nucleus neurons in heart failure. Am J Physiol Heart Circ Physiol. 2002;283 (1):H423–433. doi: 10.1152/ajpheart.00685.2001.

11. Xue B, Zhang Y, Johnson AK. Interactions of the Brain Renin-Angiotensin-System (RAS) and Inflammation in the Sensitization of Hypertension. Front Neurosci. 2020;14:650. doi: 10.3389/fnins.2020.00650.

12. Coble JP, Grobe JL, Johnson AK, Sigmund CD. Mechanisms of brain renin angiotensin system-induced drinking and blood pressure: importance of the subfornical organ. Am J Physiol Regul Integr Comp Physiol. 2015;308 (4):R238–249. doi: 10.1152/ajpregu.00486.2014.

13. Souza LAC, Worker CJ, Li W, et al. (Pro)renin receptor knockdown in the paraventricular nucleus of the hypothalamus attenuates hypertension development and AT(1) receptor-mediated calcium events. Am J Physiol Heart Circ Physiol. 2019;316 (6):H1389–H1405. doi: 10.1152/ajpheart.00780.2018.

14. Fountain JH, Kaur J, Lappin SL. Physiology, Renin Angiotensin System. In: StatPearls. Treasure Island (FL)2025.

15. Souza LAC, Earley YF. (Pro)renin Receptor and Blood Pressure Regulation: A Focus on the Central Nervous System. Curr Hypertens Rev. 2022;18 (2):101–116. doi: 10.2174/1570162X20666220127105655.

16. Xia H, de Queiroz TM, Sriramula S, Feng Y, Johnson T, Mungrue IN, Lazartigues E. Brain ACE2 overexpression reduces DOCA-salt hypertension independently of endoplasmic reticulum stress. Am J Physiol Regul Integr Comp Physiol. 2015;308 (5):R370–378. doi: 10.1152/ajpregu.00366.2014.

17. Worker CJ, Li W, Feng CY, et al. The neuronal (pro)renin receptor and astrocyte inflammation in the central regulation of blood pressure and blood glucose in mice fed a high-fat diet. Am J Physiol Endocrinol Metab. 2020;318 (5):E765–E778. doi: 10.1152/ajpendo.00406.2019.

18. de Kloet AD, Pati D, Wang L, et al. Angiotensin type 1a receptors in the paraventricular nucleus of the hypothalamus protect against diet-induced obesity. J Neurosci. 2013;33 (11):4825–4833. doi: 10.1523/JNEUROSCI.3806-12.2013.

19. Kang YM, Ma Y, Zheng JP, Elks C, Sriramula S, Yang ZM, Francis J. Brain nuclear factor-kappa B activation contributes to neurohumoral excitation in angiotensin II-induced hypertension. Cardiovasc Res. 2009;82 (3):503–512. doi: 10.1093/cvr/cvp073.

20. Feng Earley Y, Pan S, Verma H, Zheng H, Plata AA, Zubcevic J, Leenen FHH. Central nervous system mechanisms of salt-sensitive hypertension. Physiol Rev. 2025;105 (4):1989–2032. doi: 10.1152/physrev.00035.2024.

21. Oh JM, An M, Son DS, Choi J, Cho YB, Yoo CE, Park WY. Comparison of cell type distribution between single-cell and single-nucleus RNA sequencing: enrichment of adherent cell types in single-nucleus RNA sequencing. Exp Mol Med. 2022;54 (12):2128–2134. doi: 10.1038/s12276-022-00892-z.

22. Waag R, Bohacek J. Single-Nucleus RNA-Sequencing in Brain Tissue. Curr Protoc. 2023;3 (11):e919. doi: 10.1002/cpz1.919.

23. Minati MA, Fages A, Dauguet N, Zhu J, Jacquemin P. Optimized nucleus isolation protocol from frozen mouse tissues for single nucleus RNA sequencing application. Front Cell Dev Biol. 2023;11:1243863. doi: 10.3389/fcell.2023.1243863.

24. Pan S, L ACS, Worker CJ, et al. (Pro)renin receptor signaling in hypothalamic tyrosine hydroxylase neurons is required for obesity-associated glucose metabolic impairment. JCI Insight. 2024;9 (6). doi: 10.1172/jci.insight.174294.

25. Gayban AJB, Souza LAC, Cooper SG, Regalado E, Kleemann R, Feng Earley Y. (Pro)Renin Receptor Antagonism Attenuates High-Fat-Diet-Induced Hepatic Steatosis. Biomolecules. 2023;13 (1). doi: 10.3390/biom13010142.

26. Cooper SG, Souza LAC, Worker CJ, Gayban AJB, Buller S, Satou R, Feng Earley Y. Renin-a in the Subfornical Organ Plays a Critical Role in the Maintenance of Salt-Sensitive Hypertension. Biomolecules. 2022;12 (9). doi: 10.3390/biom12091169.

27. Li W, Peng H, Mehaffey EP, et al. Neuron-specific (pro)renin receptor knockout prevents the development of salt-sensitive hypertension. Hypertension. 2014;63 (2):316–323. doi: 10.1161/HYPERTENSIONAHA.113.02041.

28. Hao Y, Stuart T, Kowalski MH, et al. Dictionary learning for integrative, multimodal and scalable single-cell analysis. Nature biotechnology. 2024;42 (2):293–304.

29. Hao Y, Hao S, Andersen-Nissen E, et al. Integrated analysis of multimodal single-cell data. Cell. 2021;184 (13):3573–3587.

30. Stuart T, Butler A, Hoffman P, et al. Comprehensive integration of single-cell data. cell. 2019;177 (7):1888–1902.

31. Butler A, Hoffman P, Smibert P, Papalexi E, Satija R. Integrating single-cell transcriptomic data across different conditions, technologies, and species. Nature biotechnology. 2018;36 (5):411–420.

32. Hafemeister C, Satija R. Normalization and variance stabilization of single-cell RNA-seq data using regularized negative binomial regression. Genome biology. 2019;20 (1):296.

33. Abdi H, Williams LJ. Principal component analysis. Wiley interdisciplinary reviews: computational statistics. 2010;2 (4):433–459.

34. McInnes L, Healy J, Melville J. Umap: Uniform manifold approximation and projection for dimension reduction. arXiv preprint arXiv:180203426. 2018.

35. Van der Maaten L, Hinton G. Visualizing data using t-SNE. Journal of machine learning research. 2008;9 (11).

36. Yao Z, Liu H, Xie F, et al. An integrated transcriptomic and epigenomic atlas of mouse primary motor cortex cell types. Biorxiv. 2020:2020-2002.

37. Bigalke JA, Gao H, Chen QH, Shan Z. Activation of Orexin 1 Receptors in the Paraventricular Nucleus Contributes to the Development of Deoxycorticosterone Acetate-Salt Hypertension Through Regulation of Vasopressin. Front Physiol. 2021;12:641331. doi: 10.3389/fphys.2021.641331.

38. Li DP, Pan HL. Glutamatergic inputs in the hypothalamic paraventricular nucleus maintain sympathetic vasomotor tone in hypertension. Hypertension. 2007;49 (4):916–925. doi: 10.1161/01.HYP.0000259666.99449.74.

39. Pietranera L, Saravia FE, McEwen BS, Lucas LL, Johnson AK, De Nicola AF. Changes in Fos expression in various brain regions during deoxycorticosterone acetate treatment: relation to salt appetite, vasopressin mRNA and the mineralocorticoid receptor. Neuroendocrinology. 2001;74 (6):396–406. doi: 10.1159/000054706.

40. Genain C, Bouhnik J, Tewksbury D, Corvol P, Menard J. Characterization of plasma and cerebrospinal fluid human angiotensinogen and des-angiotensin I- angiotensinogen by direct radioimmunoassay. J Clin Endocrinol Metab. 1984;59 (3):478–484. doi: 10.1210/jcem-59-3-478.

41. Sherrod M, Liu X, Zhang X, Sigmund CD. Nuclear localization of angiotensinogen in astrocytes. Am J Physiol Regul Integr Comp Physiol. 2005;288 (2):R539–546. doi: 10.1152/ajpregu.00594.2004.

42. Stornetta RL, Hawelu-Johnson CL, Guyenet PG, Lynch KR. Astrocytes synthesize angiotensinogen in brain. Science. 1988;242 (4884):1444–1446. doi: 10.1126/science.3201232.

43. Elsaafien K, Kirchner MK, Mohammed M, et al. Identification of Novel Cross-Talk between the Neuroendocrine and Autonomic Stress Axes Controlling Blood Pressure. J Neurosci. 2021;41 (21):4641–4657. doi: 10.1523/JNEUROSCI.0251-21.2021.

44. Colmers PLW, Bains JS. Balancing tonic and phasic inhibition in hypothalamic corticotropin-releasing hormone neurons. J Physiol. 2018;596 (10):1919–1929. doi: 10.1113/JP275588.

45. Holbein WW, Blackburn MB, Andrade MA, Toney GM. Burst patterning of hypothalamic paraventricular nucleus-driven sympathetic nerve activity in ANG II- salt hypertension. Am J Physiol Heart Circ Physiol. 2018;314 (3):H530–H541. doi: 10.1152/ajpheart.00560.2017.

46. Sladek CD, Michelini LC, Stachenfeld NS, Stern JE, Urban JH. Endocrine-Autonomic Linkages. Compr Physiol. 2015;5 (3):1281–1323. doi: 10.1002/cphy.c140028.

47. Lenkei Z, Palkovits M, Corvol P, Llorens-Cortes C. Expression of angiotensin type-1 (AT1) and type-2 (AT2) receptor mRNAs in the adult rat brain: a functional neuroanatomical review. Front Neuroendocrinol. 1997;18 (4):383–439. doi: 10.1006/frne.1997.0155.

48. de Kloet AD, Wang L, Ludin JA, et al. Reporter mouse strain provides a novel look at angiotensin type-2 receptor distribution in the central nervous system. Brain Struct Funct. 2016;221 (2):891–912. doi: 10.1007/s00429-014-0943-1.

49. Lenkei Z, Palkovits M, Corvol P, Llorens-Cortes C. Distribution of angiotensin II type- 2 receptor (AT2) mRNA expression in the adult rat brain. J Comp Neurol. 1996;373 (3):322–339. doi: 10.1002/(SICI)1096-9861(19960923)373:3<322::AID-CNE2>3.0.CO;2-4.

50. Coleman CG, Anrather J, Iadecola C, Pickel VM. Angiotensin II type 2 receptors have a major somatodendritic distribution in vasopressin-containing neurons in the mouse hypothalamic paraventricular nucleus. Neuroscience. 2009;163 (1):129–142. doi: 10.1016/j.neuroscience.2009.06.032.

51. Reagan LP, Flanagan-Cato LM, Yee DK, Ma LY, Sakai RR, Fluharty SJ. Immunohistochemical mapping of angiotensin type 2 (AT2) receptors in rat brain. Brain Res. 1994;662 (1-2):45–59. doi: 10.1016/0006-8993(94)90794-3.

52. Fekete EM, Gomez J, Ghobrial M, et al. Definitive Evidence for the Identification and Function of Renin-Expressing Cholinergic Neurons in the Nucleus Ambiguus. Hypertension. 2025;82 (2):282–292. doi: 10.1161/HYPERTENSIONAHA.124.23740.

53. Hersh LB, Aboukhair N, Watson S. Immunohistochemical localization of aminopeptidase M in rat brain and periphery: relationship of enzyme localization and enkephalin metabolism. Peptides. 1987;8 (3):523–532. doi: 10.1016/0196-9781(87)90019-2.

54. Solhonne B, Gros C, Pollard H, Schwartz JC. Major localization of aminopeptidase M in rat brain microvessels. Neuroscience. 1987;22 (1):225–232. doi: 10.1016/0306-4522(87)90212-0.

55. Lochard N, Thibault G, Silversides DW, Touyz RM, Reudelhuber TL. Chronic production of angiotensin IV in the brain leads to hypertension that is reversible with an angiotensin II AT1 receptor antagonist. Circ Res. 2004;94 (11):1451–1457. doi: 10.1161/01.RES.0000130654.56599.40.

56. Jin X, Kim WB, Kim MN, et al. Oestrogen inhibits salt-dependent hypertension by suppressing GABAergic excitation in magnocellular AVP neurons. Cardiovasc Res. 2021;117 (10):2263–2274. doi: 10.1093/cvr/cvaa271.

57. Pitra S, Worker CJ, Feng Y, Stern JE. Exacerbated effects of prorenin on hypothalamic magnocellular neuronal activity and vasopressin plasma levels during salt-sensitive hypertension. Am J Physiol Heart Circ Physiol. 2019;317 (3):H496–H504. doi: 10.1152/ajpheart.00063.2019.

58. Pitra S, Feng Y, Stern JE. Mechanisms underlying prorenin actions on hypothalamic neurons implicated in cardiometabolic control. Mol Metab. 2016;5 (10):858–868. doi: 10.1016/j.molmet.2016.07.010.

59. Haywood JR, Mifflin SW, Craig T, Calderon A, Hensler JG, Hinojosa-Laborde C. gamma-Aminobutyric acid (GABA)--A function and binding in the paraventricular nucleus of the hypothalamus in chronic renal-wrap hypertension. Hypertension. 2001;37 (2 Pt 2):614–618. doi: 10.1161/01.hyp.37.2.614.

60. Martin DS, Haywood JR. Reduced GABA inhibition of sympathetic function in renal- wrapped hypertensive rats. Am J Physiol. 1998;275 (5):R1523–1529. doi: 10.1152/ajpregu.1998.275.5.R1523.

61. Li DP, Pan HL. Plasticity of GABAergic control of hypothalamic presympathetic neurons in hypertension. Am J Physiol Heart Circ Physiol. 2006;290 (3):H1110–1119. doi: 10.1152/ajpheart.00788.2005.

62. Basting T, Xu J, Mukerjee S, Epling J, Fuchs R, Sriramula S, Lazartigues E. Glutamatergic neurons of the paraventricular nucleus are critical contributors to the development of neurogenic hypertension. J Physiol. 2018;596 (24):6235–6248. doi: 10.1113/JP276229.

63. Li W, Peng H, Cao T, et al. Brain-targeted (pro)renin receptor knockdown attenuates angiotensin II-dependent hypertension. Hypertension. 2012;59 (6):1188–1194. doi: 10.1161/HYPERTENSIONAHA.111.190108.

64. Brinkmann BG, Agarwal A, Sereda MW, et al. Neuregulin-1/ErbB signaling serves distinct functions in myelination of the peripheral and central nervous system. Neuron. 2008;59 (4):581–595. doi: 10.1016/j.neuron.2008.06.028.

65. Franklin RJ, Ffrench-Constant C. Remyelination in the CNS: from biology to therapy. Nat Rev Neurosci. 2008;9 (11):839–855. doi: 10.1038/nrn2480.

66. Siffrin V, Vogt J, Radbruch H, Nitsch R, Zipp F. Multiple sclerosis - candidate mechanisms underlying CNS atrophy. Trends Neurosci. 2010;33 (4):202–210. doi: 10.1016/j.tins.2010.01.002.

67. Langley MR, Yoon H, Kim HN, et al. High fat diet consumption results in mitochondrial dysfunction, oxidative stress, and oligodendrocyte loss in the central nervous system. Biochim Biophys Acta Mol Basis Dis. 2020;1866 (3):165630. doi: 10.1016/j.bbadis.2019.165630.

68. Maiuolo J, Gliozzi M, Musolino V, et al. Environmental and Nutritional "Stressors" and Oligodendrocyte Dysfunction: Role of Mitochondrial and Endoplasmatic Reticulum Impairment. Biomedicines. 2020;8 (12). doi: 10.3390/biomedicines8120553.

69. Mazier W, Saucisse N, Simon V, Cannich A, Marsicano G, Massa F, Cota D. mTORC1 and CB1 receptor signaling regulate excitatory glutamatergic inputs onto the hypothalamic paraventricular nucleus in response to energy availability. Mol Metab. 2019;28:151–159. doi: 10.1016/j.molmet.2019.08.005.

70. Clyburn C, Travagli RA, Browning KN. Acute high-fat diet upregulates glutamatergic signaling in the dorsal motor nucleus of the vagus. Am J Physiol Gastrointest Liver Physiol. 2018;314 (5):G623–G634. doi: 10.1152/ajpgi.00395.2017.

71. Zhu C, Xu Y, Jiang Z, et al. Disrupted hypothalamic CRH neuron responsiveness contributes to diet-induced obesity. EMBO Rep. 2020;21 (7):e49210. doi: 10.15252/embr.201949210.

72. Szczepanska-Sadowska E, Wsol A, Cudnoch-Jedrzejewska A, Czarzasta K, Zera T. Multiple Aspects of Inappropriate Action of Renin-Angiotensin, Vasopressin, and Oxytocin Systems in Neuropsychiatric and Neurodegenerative Diseases. J Clin Med. 2022;11 (4). doi: 10.3390/jcm11040908.

73. Grobe JL, Buehrer BA, Hilzendeger AM, Liu X, Davis DR, Xu D, Sigmund CD. Angiotensinergic signaling in the brain mediates metabolic effects of deoxycorticosterone (DOCA)-salt in C57 mice. Hypertension. 2011;57 (3):600–607. doi: 10.1161/HYPERTENSIONAHA.110.165829.

74. Basting T, Lazartigues E. DOCA-Salt Hypertension: an Update. Curr Hypertens Rep. 2017;19 (4):32. doi: 10.1007/s11906-017-0731-4.

75. Hilzendeger AM, Cassell MD, Davis DR, Stauss HM, Mark AL, Grobe JL, Sigmund CD. Angiotensin type 1a receptors in the subfornical organ are required for deoxycorticosterone acetate-salt hypertension. Hypertension. 2013;61 (3):716–722. doi: 10.1161/HYPERTENSIONAHA.111.00356.

76. Mendoza A, Lazartigues E. The compensatory renin-angiotensin system in the central regulation of arterial pressure: new avenues and new challenges. Ther Adv Cardiovasc Dis. 2015;9 (4):201–208. doi: 10.1177/1753944715578056.

77. Soto JE, Burnett CML, Ten Eyck P, Abel ED, Grobe JL. Comparison of the Effects of High-Fat Diet on Energy Flux in Mice Using Two Multiplexed Metabolic Phenotyping Systems. Obesity (Silver Spring). 2019;27 (5):793–802. doi: 10.1002/oby.22441.

78. do Carmo JM, da Silva AA, Moak SP, Browning JR, Dai X, Hall JE. Increased sleep time and reduced energy expenditure contribute to obesity after ovariectomy and a high fat diet. Life Sci. 2018;212:119–128. doi: 10.1016/j.lfs.2018.09.034.

79. Tramunt B, Smati S, Grandgeorge N, Lenfant F, Arnal JF, Montagner A, Gourdy P. Sex differences in metabolic regulation and diabetes susceptibility. Diabetologia. 2020;63 (3):453–461. doi: 10.1007/s00125-019-05040-3.

80. Prajapati C, Koivumaki J, Pekkanen-Mattila M, Aalto-Setala K. Sex differences in heart: from basics to clinics. Eur J Med Res. 2022;27 (1):241. doi: 10.1186/s40001-022-00880-z.

